# Circulatory proteins shape microglia state and boost phagocytosis

**DOI:** 10.1101/2024.09.30.615861

**Authors:** Nannan Lu, Patricia Moran-Losada, Oliver Hahn, Aryaman Saksena, Emma Tapp, Jean Paul Chadarevian, Wentao Dong, Sophia M. Shi, Steven R. Shuken, Ian Guldner, Wenshu Zeng, Ning-Sum To, Pui Shuen Wong, Jonathan Hasselmann, Hayk Davtyan, Jerry Sun, Lulin Li, Jian Luo, Andrew C. Yang, Qingyun Li, Tom H. Cheung, Monther Abu-Remaileh, Mathew Blurton-Jones, Tony Wyss-Coray

**Author notes:** These authors contributed equally (Patricia Moran-Losada, Oliver Hahn, Aryaman Saksena). Lead contract (Tony Wyss-Coray).

## Abstract

Microglia, the brain’s immune cells, are highly responsive to their local environment. Given that circulatory proteins can enter the brain, we asked whether microglia are responsive to such proteins. Here, we identify a stable population of microglia specialized to take up circulatory proteins in a region-specific manner under physiological conditions; human hematopoietic stem cell-derived microglia replacing endogenous microglia in chimeric mice show similar regional specialization. Plasma-positive microglia are characterized by prominent expression of genes related to innate immunity and antigen presentation and exhibit high metabolic and phagocytic activity. This activity is dependent, in part, on microglial uptake and accumulation of circulatory Apolipoprotein AI (ApoA-I). Our findings thus identify a new model of communication between brain and periphery through specialized microglia.

## Introduction

Interorgan communication at the whole-body level and intercellular communication within the brain are crucial for proper nervous system function in health and disease^1–4^. Soluble factors in plasma, including proteins, lipoprotein particles, and metabolites, play critical roles as messengers for interorgan communication throughout the body and for regulating brain functions^5,6^. For example, β2-microglobulin (B2M), a major histocompatibility complex class I (MHC-I) component in blood, acts as an endogenous NMDAR antagonist, and has been shown to contribute to synaptic deficits in Down Syndrome^7^. Apolipoprotein J (ApoJ), also known as clusterin, is upregulated in plasma following physical exercise, reducing brain inflammation, and enhancing cognitive memory^8^. N-lactoyl-phenylalanine (Lac-Phe), a metabolite induced by physical exercise in the blood, suppresses food intake and mitigates obesity^9^.

Recent studies reveal the potential for blood-based interventions, such as heterochronic parabiosis or systemic administration of young blood plasma component into aged mice^10–13^, to rejuvenate cognitive function in aged brains by reducing neuronal and glial inflammation, remodeling vasculature, and increasing neurogenesis. Plasma protein factors likely contribute to these brain rejuvenating effects through multiple pathways. Some proteins are actively or passively transported across the BBB into the brain parenchyma, while others may penetrate regions with a permeable blood-brain barrier such as circumventricular organs and directly interact with other brain cell types. Additionally, certain proteins that are unable to penetrate the brain may still impact brain functions indirectly by affecting the endothelium or platelet factors, as exemplified by ApoE4^14^ and klotho^15^. Our previous work utilizing labeled whole-blood plasma proteome in young mice revealed a substantial amount of protein taken up by brain endothelial cells, neurons, and glia in the brain parenchyma^16^. This suggests that plasma proteins readily permeate the healthy brain parenchyma. Yet, to what extent and how plasma proteins modulate the function of brain cells within the parenchyma under physiological conditions remains largely unknown.

Microglia, the resident macrophages of the brain, play critical roles in maintaining central nervous system (CNS) homeostasis and plasticity^17–19^. These specialized cells continuously surveil the brain and are highly responsive to local environmental cues. They sense the surrounding microenvironment – including neurotransmitters, apoptotic cells, extracellular debris, and amyloid-beta plaques^20,21^ – via surface receptors and transporters. This sensing modulates their behavior producing changes in morphology, transcriptional state, metabolic activity, motility, and function. Microglia also rapidly adapt to their physical location and engage in reciprocal interactions with neighboring cells and structures. In addition, Microglia are highly responsive to long distance signals originating outside of the brain, such as diet-produced fatty acids and products from the gut microbiota^22–24^. Notably, maternal omega-3 fatty acid deficiency increases microglia-mediated phagocytosis of synaptic elements, alters neuronal morphology, and impairs cognitive performance in offspring^25^. Through the microbiota-gut-brain axis, short-chain fatty acids derived from gut microbiota act as key regulators, influencing microglial function and maturation in health and disease^22,24^. Disruption of the microbiome perturbs microglial homeostasis, resulting in impaired innate immune responses. Thus, microglia can serve as a sensitive hub and possible integrator of systemic signals in the brain. After sensing signals, they further transmit their responses to regulate neuronal activity, pruning synapses, and modulating behaviors^26,27^. Building upon our previous observations showing that plasma proteins readily permeate the healthy brain parenchyma, we hypothesize that microglia actively respond to blood-borne factors. In this study, by labeling and tracing plasma proteins *in vivo*, we demonstrate that circulatory factors directly regulate microglia function in healthy brains. Interestingly, plasma-positive microglia exhibit increased innate immune response, higher metabolic activity, and phagocytic capacity than plasma-negative microglia. We also identify ApoA-I as one of the prominent plasma proteins taken up by microglia and acting as a positive regulator of microglial phagocytosis.

## Results

### Specialized microglia take up plasma proteins in a region-specific manner

To investigate the impact of circulatory factors on microglia under adult physiological conditions, we labeled the entire mouse plasma proteome with a small far-red fluorophore dye (NHS-Atto647N) using N-hydroxysuccinimide (NHS) ester chemistry as previously reported^16^. Twenty hours after intravenously injecting dye-labeled plasma proteins into recipient male mice (3-month-old), we extensively perfused them and isolated microglia from six major brain regions, including: hypothalamus (HTH), thalamus (TH), hippocampus (HP), striatum (STR), cortex (CTX), and cerebellum (CB, Figure 1A) using a cold mechanical dissociation method^28,29^. Flow cytometry analysis revealed remarkable regional heterogeneity in labeled plasma uptake by microglia within the brain, with the highest uptake observed in the hypothalamus (15.28% of total microglia), followed by the thalamus (10.02%) and hippocampus (6.08%), and much lower uptake in the cerebellum (2.48%), striatum (2.26%), and cortex (0.76%, Figure 1B-C, Figure S1A). Additionally, the mean fluorescence intensity of plasma-positive-microglia (PPM) was higher in the hypothalamus compared to the thalamus and hippocampus (Figure 1D). A similar regional heterogeneity was observed in female mice (Figure S1B), arguing against sex-dependence of the observed phenomenon. This region-specific uptake was also observed when plasma proteins were labeled with maleimide Atto550, which primarily labels thiol-containing cysteine residues (Figure S1C). No detectable plasma uptake was observed with carboxy-Atto647N, which does not effectively label proteins, suggesting our observations depend on administration of dye-labeled plasma proteins (Figure S1D). Furthermore, plasma signals in microglia were not a result of contamination during *ex vivo* cell dissociation. Plasma-positive microglia were exclusively observed in wild-type but not Ubi-GFP mice when mixing brain tissue from Ubi-GFP mice uninjected and tissue from wild-type mice injected with labeled plasma proteins during the tissue dissociation steps (Figure S1E-G). Plasma uptake in microglia increased gradually, reaching a peak at 20-40 hours after injection in both hypothalamus and thalamus (Figure S1H and I).

**Figure 1.**
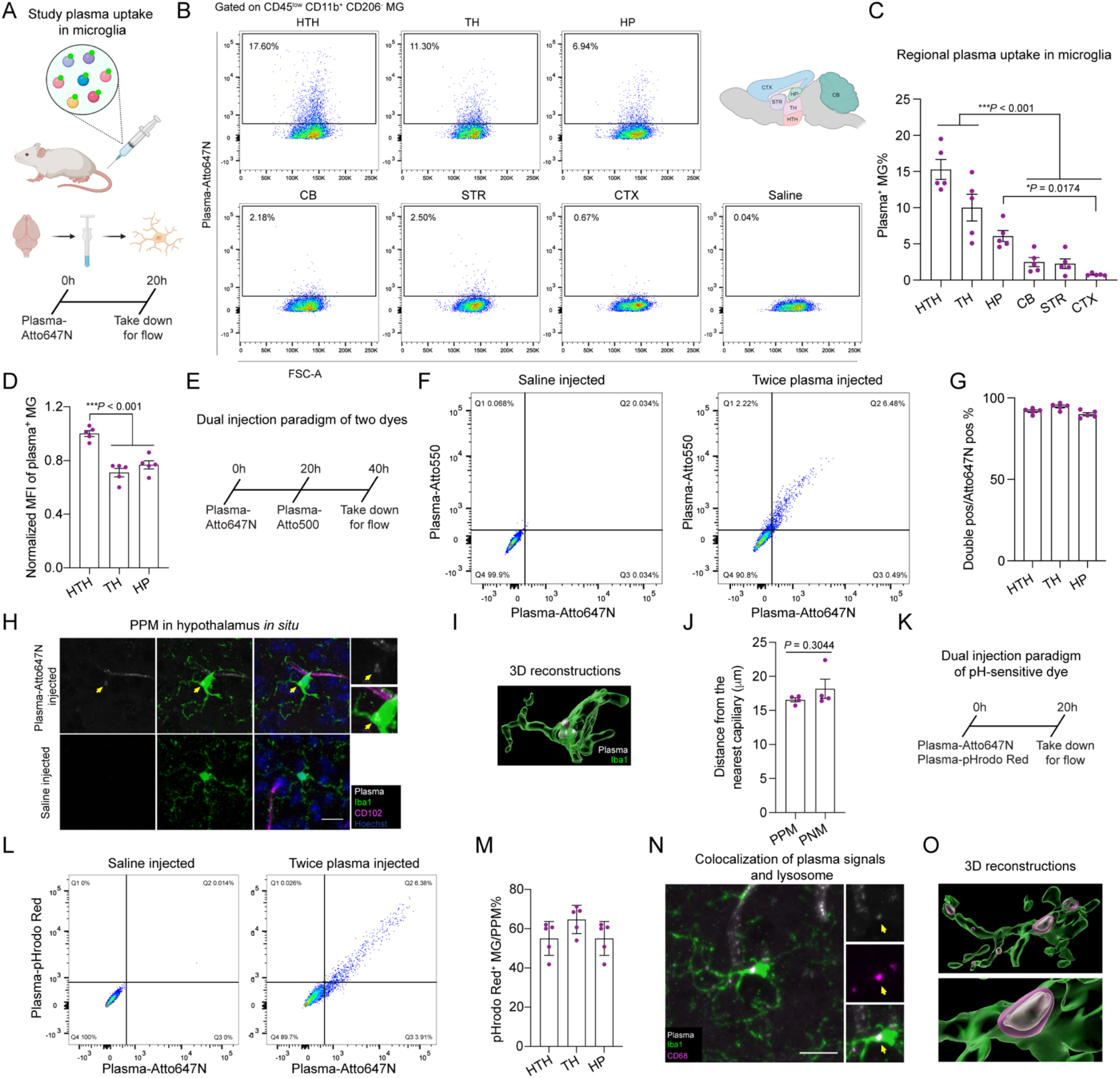
Microglia take up plasma proteins from blood circulation in a brain-region specific manner. A) Experimental scheme for measuring the uptake of NHS-Atto647N labeled plasma proteins in microglia by flow cytometry. B) Representative flow cytometry data of the same number of pre-gated microglia (CD45^low^CD11b^+^CD206^-^) in six brain regions of labeled plasma-injected mouse comparing to saline-injected mouse. The percentages of plasma-positive microglia in each region are shown. C) Quantification of the percentage of plasma-positive microglia versus total microglia in six brain regions of 3-month-old mice (n = 5 mice per group; mean ± s.e.m.; one-way ANOVA, followed by Tukey’s post hoc test). D) Normalized geometric mean fluorescence intensity (MFI) of plasma-positive microglia in three brain regions in 3-month-old mice (n = 5 mice per group; mean ± s.e.m.; one-way ANOVA, followed by Tukey’s post hoc test). E) Experimental scheme for injecting 2 different dyes (NHS-Atto647N and NHS-Atto550) labeled plasma proteins into the same mouse with 20 hours apart. F) Representative flow cytometry data of pre-gated microglia from twice plasma protein-injected mouse versus saline-injected mouse. The percentages of cells in each quadrant are shown. Q2 shows the double-positive microglia. G) Quantification of the percentage of double dye-positive microglia versus Atto647N-plasma positive microglia in three brain regions (n = 5 mice per group; mean ± s.e.m.). H) Fluorescence microscopy images of plasma uptake in microglia (Iba1^+^) from hypothalamus of 3-month-old mice. Scale bar, 15 μm. The yellow arrow indicates the Atto647N labeled plasma signals within microglia. CD102 stains brain endothelial cells. Hoechst labels cell nuclei. I) 3D reconstruction shows plasma protein signals (grey) are inside microglia (green). J) Quantification of the distances of plasma-positive microglia (PPM) and plasma-negative microglia (PNM) to the nearest capillary in hypothalamus (n = 4 mice per group; mean ± s.e.m.; unpaired two-sided *t*-test). K) Experimental scheme of injecting Atto647N-labeled plasma proteins and pHrodo-Red-labeled plasma proteins into the same mouse. Flow cytometry assays were performed 20 hours later to analyze the uptake of the labeled plasma proteins in the microglia. L) Representative flow data of pre-gated microglia from twice plasma protein-injected mouse (Atto647N and pHrodo-Red-labeled plasma) versus the saline-injected mouse. The percentages of cells in each quadrant are shown. Q2 shows the double-positive microglia. M) Quantification of the percentage of pHrodo Red dye-positive microglia versus Atto647N-positive microglia in three brain regions (n = 5 mice per group; mean ± s.e.m.). N) Subcellular localizations of plasma proteins within microglia (Iba1^+^) in hypothalamus of 3-month-old mice injected with Atto647N-labeled plasma proteins. Yellow arrow indicates that the Atto647N-labeled plasma signals are overlapping with lysosome (CD68^+^). Scale bar, 15 μm. O) 3D reconstruction shows the plasma protein signals (grey) are overlapping with lysosomes (magenta) in microglia (green). HTH, hypothalamus; TH, thalamus; HP, hippocampus; CB, cerebellum; STR, striatum; CTX, cortex. See also Figure S1.

To investigate whether PPM represent a specialized population, or any microglia can internalize plasma proteins from the periphery, we injected plasma proteins labeled with two different fluorophores (NHS-Atto647N and NHS-Atto550) into the same mouse with a twenty-hour interval between injections. Interestingly, among the NHS-Atto647N-plasma positive microglia, >95% of them were double-positive for both dyes (Figure 1E-G), indicating that, under the current plasma injection paradigm, this subset of microglia is a stable population that is specialized to take up circulatory proteins. Consistent with an overall decrease in plasma protein uptake with age^16^, we observed a 60% decrease in labeled plasma protein uptake by hypothalamic and hippocampal microglia in 24-month-old compared with 3-month-old mice, while other brain regions were unchanged (Figure S1J). These findings highlight the specificity of plasma protein uptake into microglia and demonstrate that it is age dependent.

Additionally, we were able to document the presence of PPM in hypothalamic brain slices in situ in mice (Figure 1H and I). Because PPM and plasma-negative-microglia (PNM) showed similar distances on average to the nearest capillary, we speculate that factors beyond proximity to the blood-brain barrier (BBB) regulate plasma uptake (Figure 1J). To determine the subcellular location of plasma proteins following microglial internalization, we used pHrodo red, a pH-sensitive dye, to label plasma proteins and injected them into the same mouse alongside NHS-Atto647N-labeled plasma proteins. Twenty hours later, dye double-positive microglia were detected in the hypothalamus, thalamus, and hippocampus (Figure 1K-M). The intensities between the two dyes correlated positively (Figure S1K). indicating that some plasma proteins were transported into acidic organelles, including the lysosome. Furthermore, plasma protein signals were co-localized with CD68^+^ lysosomes in hypothalamic microglia (Figure 1N and O). Collectively, our data uncover the existence of a population of microglia specialized to take up circulatory proteins under physiological conditions. We find brain region specific heterogeneity in the percentage of the PPMs and how they change with age.

### Plasma-positive microglia exhibit distinct molecular signatures compared to Plasma-negative microglia

To investigate whether regional heterogeneity of plasma protein uptake correlates with microglial transcriptome diversity, we isolated microglia from six major brain regions (hypothalamus, thalamus, hippocampus, striatum, cortex, and cerebellum) from 3-month-old mice and performed bulk RNA sequencing (RNA-seq) analysis (Figure S2A). Principal Component Analysis (PCA) reveals a significant distinction between microglia from the cerebellum and those from other brain regions, aligning with previous findings^30,31^. Microglia from the hypothalamus and thalamus exhibit an intermediate status among the six regions, while microglia from the striatum, hippocampus and cortex share the greatest similarity (Figure S2B and C). However, this pattern of transcriptomic similarity among these regions does not explain the higher plasma protein uptake in the hypothalamus and the lower uptake in the cerebellum.

To gain deeper insight into the transcriptional characteristics of PPM and PNM from the young brain, we isolated these microglia (CD11b^+^CD45^low^CD206^-^) from hypothalamus of mice (3-month-old) injected with Atto647N-labeled plasma proteins based on the intensity of Atto647N fluorescence using fluorescence-activated cell sorting (FACS) and analyzed them by bulk RNA-seq (Figure 2A). Importantly, this region was chosen due to the high percentage and intensity of plasma protein uptake by microglia within the hypothalamus (Figure 1C and D). In addition, we use cold mechanical tissue homogenization to maintain microglia in a non-active state and minimize bias towards an activated proinflammatory signature^32,33^. Confocal images of *ex vivo* dissociated, and FACS-sorted microglia confirmed the presence of plasma protein signals within PPM (Figure 2B). A pronounced separation between PPM and PNM at the transcriptomic level in the young healthy brains was observed from the PCA plot (Figure 2C; Table S1). Expression analysis identified 576 differentially expressed genes (DEGs) significantly upregulated in PPM, while 579 DEGs were downregulated (Figure 2D). Pathway analysis of differentially expressed genes revealed several key pathways. Notably, pathways associated with the innate immune response (*Apoe, Axl, Csf1r, C1qb, Ifitm3*), chemokine-mediated signaling, chemotaxis (*Ccl5, Ccl6, Ccl9, Lgals3*), endocytosis (*Axl, Cd63, Fcer1g*), and antigen presentation (*B2m, H2-D1*) were upregulated in PPM as compared to PNM (Figure 2E; Table S2).

**Figure 2.**
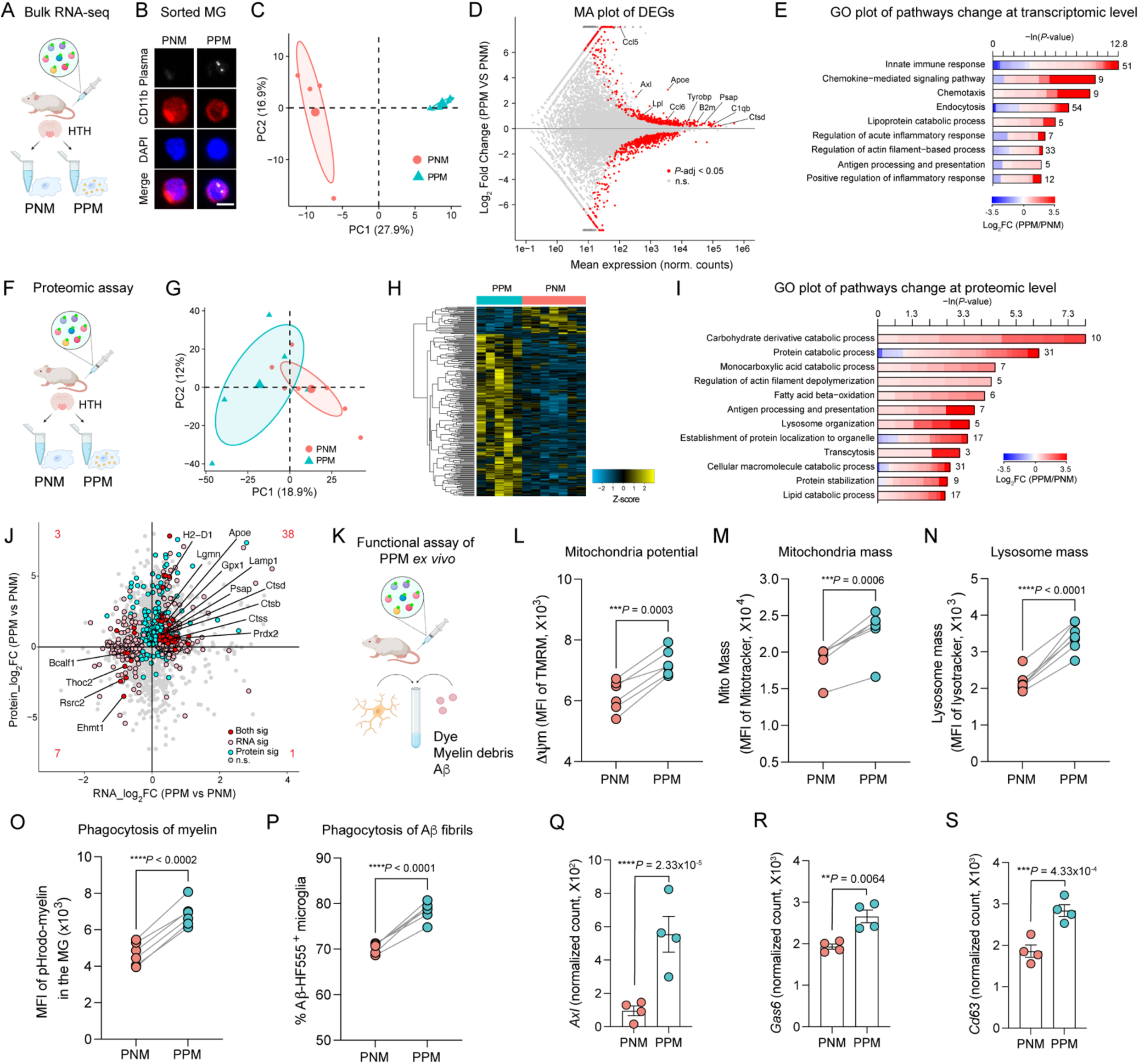
Increased metabolic activity and phagocytosis in plasma-positive microglia compared to plasma-negative microglia from young mice. A) Experimental scheme for sorting plasma-positive microglia (PPM) and plasma-negative microglia (PNM) from hypothalamus (HTH) of 3-month-old young mice 20 hours after injecting NHS-Atto647N labeled plasma proteins. The sorted microglia were then used for bulk RNA-seq. B) Representative images of FACS-sorted PPM and PNM from young mice (scale bar, 5 μm). Plasma signals were observed inside microglia. C) Principal component analysis (PCA) plot of bulk RNA-seq transcriptomes of PPM and PNM isolated from the hypothalamus of young mice injected with NHS-Atto647N labeled plasma proteins (n = 4 mice per group). D) MA-plot representing log_2_-transformed gene expression fold change in PPM/PNM versus the mean expression on a logarithmic scale. Genes showing significant differential expression between two groups are marked in red (n = 1155 genes; *P*-adjust < 0.05, Wald test, followed by FDR/Benjamini– Hochberg test). E) Representative Gene Ontology (GO) enrichment categories of PPM versus PNM are shown. The lengths of the bars represent the negative natural logarithm (ln) of transformed *P*-value obtained using a two-sided Fisher’s exact test. The colors indicate gene-wise log_2_ fold-changes between PPM versus PNM. The numbers beside the bars indicate the number of differential expressed genes (DEGs) in that specific GO category. F) Experimental scheme for sorting PPM and PNM from the hypothalamus (HTH) of 3-month-old young mice 20h after injecting NHS-Atto647N labeled plasma proteins. The sorted microglia were then subjected to whole cell proteomic assay. G) PCA plot of proteome of PPM and PNM isolated from the hypothalamus of young mice injected with NHS-Atto647N labeled plasma proteins (n = 7 and n = 5 biological replicates for PNM and PPM, respectively). H) Heatmap showing the significantly differentially enriched proteins between PPM and PNM. The protein abundance is z-score transformed (*P* < 0.05, two-sided *t*-test) I) Representative GO enrichment categories of PPM versus PNM at protein level are shown. The lengths of the bars represent negative ln of transformed *P*-value using two-sided Fisher’s exact test. The colors indicate protein-wise log_2_ fold-changes between PPM versus PNM. The numbers beside the bars indicate differential expressed proteins in that specific GO category. J) Scatter plot of the log_2_ fold change of RNA and protein between PPM and PNM. Red dots indicate genes that change significantly at both transcriptome and proteome level. The number of significantly differentially expressed genes at both RNA and protein levels in each quadrant is indicated in red. K) Experimental scheme to evaluate the functional differences between PPM and PNM using *ex vivo* dye incubation and phagocytosis assays. L-N) Quantification of TMRM (L), MitoTracker Green (M), and LysoTracker Green (N) staining in PPM and PNM by flow cytometry (n= 6 mice per group for TMRM and MitoTracker; n= 7 mice per group for LysoTracker; paired two-sided *t*-test). O) Geometric mean fluorescence intensity (MFI) of *ex vivo* microglial phagocytosis of pHrodo-myelin debris in PPM versus PNM (n= 6 mice per group, paired two-sided *t*-test). P) Percentage of *ex vivo* microglial phagocytosis of Aβ fibrils (Aβ-HF555) in PPM versus PNM (n= 6 mice per group; paired two-sided *t*-test). Q-S) Normalized count of phagocytosis related genes *Axl* (Q), *Gas6* (R), *CD63* (S) from the bulk transcriptome comparing PPM versus PNM (n= 4 mice per group; mean ± s.e.m.; *P* values determined by DESeq2). Both sig, both significant; RNA sig, only RNA level significant; Protein sig, only protein level significant; n.s., not significant. See also Figure S2, Table S1-S6.

Since changes at the transcriptomic and proteomic levels may not always be directly correlated, we also investigated the proteomic profile of sorted PPM and PNM from young mice and performed whole-cell proteomic analysis using timsTOF mass spectrometry (Figure 2F). On average, 2,274 proteins were detected per sample, obtained from the sorting of 12,000 microglia per sample. PCA analysis effectively distinguished PPM from PNM (Figure 2G; Table S3), revealing that 220 proteins are significantly enriched in PPM while 37 proteins are notably down-regulated in PPM, compared with PNM (Figure 2H). Differentially enriched proteins in PPM are involved in the catabolism of carbohydrate derivatives, proteins, and monocarboxylic acids, and lipids as well as in fatty acid beta-oxidation, indicating higher metabolic activity in these cells (Figure 2I; Table S4). Consistent with transcriptomic changes (Figure 2E), antigen processing and presentation are also increased in PPM compared with PNM. Interestingly, we identified 38 genes that were increased and 7 genes that were decreased at both RNA and protein levels, including up-regulated genes involved in lysosome function (*Psap, Lamp1, Ctsd, Ctsb, and Ctss*), antigen presentation (*H2-D1*) and antioxidant pathway (*Gpx1, Prdx2*), and down-regulated genes involved in transcriptional repressor (*Bclaf1* and *Ehmt1*, Figure 2J).

Given that many cellular catabolic processes are increased in PPM, we further conducted metabolomic and lipidomic profiling using liquid chromatography–mass spectrometry (LC–MS) to compare PPM and PNM from young mice (Figure S2D). Notably, butyryl-carnitine was significantly increased in PPM, along with a trend towards an increase in two other carnitine related metabolites, acetyl-carnitine, and carnitine (Figure S2E; Table S5). These metabolites are associated with transporting long-chain fatty acids into mitochondria for oxidation and energy production^34^, suggesting higher energy metabolic activity in PPM. Lipidomic profiling of 185 lipid species from 50,000 sorted microglia per replicate revealed a significant increase of Coenzyme Q9 (Co(Q9)) and Coenzyme Q10 (Co(Q10)) in PPM (Figure S2F; Table S6). Co(Q10) not only serves as an essential redox-active lipid in the mitochondrial electron transport chain, facilitating electron relay, but also functions as a cofactor for biosynthetic and catabolic reactions^35^, thus supporting the notion of elevated mitochondria metabolic activity in PPM. Collectively, our multi-omics profiling data indicate that PPM exhibit a distinct molecular signature, characterized by upregulation of the innate immune response, antigen presentation, and enhanced catabolic and metabolic processes.

### Plasma-positive microglia exhibit higher metabolic activity and phagocytic capacity

To further determine whether these molecular changes in microglia correspond to their functional status, we conducted *ex vivo* functional profiling of hypothalamic microglia (Figure 2K). We found that PPM display higher mitochondrial activity, as indicated by the mitochondrial potential-sensitive dye TMRM^36^ (Figure 2L, Figure S2G). Additionally, PPM exhibited greater mitochondria mass, as assessed using mitoTracker staining, and showed increased lysosomal mass, determined by using lysoTracker, compared to PNM from the same mouse (Figure 2M and N, Figure S2H and I). Notably, elevated levels of the mitochondrial proteins TOM20 and COXIV were also observed in PPM (Figure S2J and K). Furthermore, when *ex vivo* microglia were incubated with pH-sensitive fluorescent myelin (phrodo-myelin) or β-Amyloid-HiLyte Fluor 555 aggregates (Aβ-HF555), PPM exhibited a significantly greater phagocytic capacity than PNM (Figure 2O and P, Figure S2L and M). This enhanced phagocytic capacity was reflected by an increase in expression levels of genes associated with phagocytosis, including *Axl*, *Gas6* and *CD63*, as observed in our bulk RNA-seq data (Figure 2Q-S). PPM also exhibited a higher level of surface CD63, as shown by antibody staining (Figure S2N). Taken together, these findings suggest that hypothalamic PPM possess a distinctive molecular signature associated with elevated metabolic activity and enhanced phagocytic functions within the healthy young brain.

### Transcriptional analysis suggests the presence of a plasma-positive microglia state in human brains

So far, our definition of plasma-positive microglia is based on the positive signals of the fluorescent tag used to label and trace plasma proteins *in vivo*. To determine if plasma-positive microglia exist in both mouse and human brains under naïve conditions, we first generated a plasma-positive gene signature by selecting the top 96 upregulated genes in PPM compared to PNM from our bulk RNA-seq analysis (Figure 2A-E, Table S7). We then applied this signature to single-cell datasets by calculating the expression of signature genes as a single plasma score for each cell (see method “scRNA-seq data analysis”)^37,38^.

Next, to validate if this plasma score can distinguish PPM from PNM, we generated an in-house droplet-based single-cell RNA-seq dataset by sorting PPM and PNM from the hypothalamus of young mice injected with Atto647N-labeled plasma (Figure 3A). Following the removal of low-quality cells and confirmation of low *ex vivo* activation score^39^ and high microglia identity score^40^ (Figure S3A-E), our dataset consisted of 17,512 high-quality microglia. Unsupervised clustering and data visualization by uniform manifold approximation and projection (UMAP) identified five distinct microglial clusters: homeostatic 1, homeostatic 2, Apoe^high^, Interferon and proliferative, as reported previously in healthy mouse brain^41^ (Figure 3B, Figure S3F). Intriguingly, the ratio of PPM/PNM was higher in the Interferon cluster (2.35-fold) compared to other clusters (Figure 3C). The Interferon cluster microglia have been identified in multiple developmental and disease models, and a recent study identified that type-I interferon-responsive microglia have a critical role in phagocytosing unwanted neuron and circuit wiring during brain development^27^. Applying the plasma score to the current dataset revealed a significantly higher plasma score in PPM compared to PNM (Figure 3D). A random selection of 100 genes from the same bulk RNA-seq dataset (Figure 2A-E, Table S7) showed no difference in the score comparison between PPM and PNM (Figure S3G-I), indicating that our plasma score is specific to the plasma uptake in microglia.

**Figure 3.**
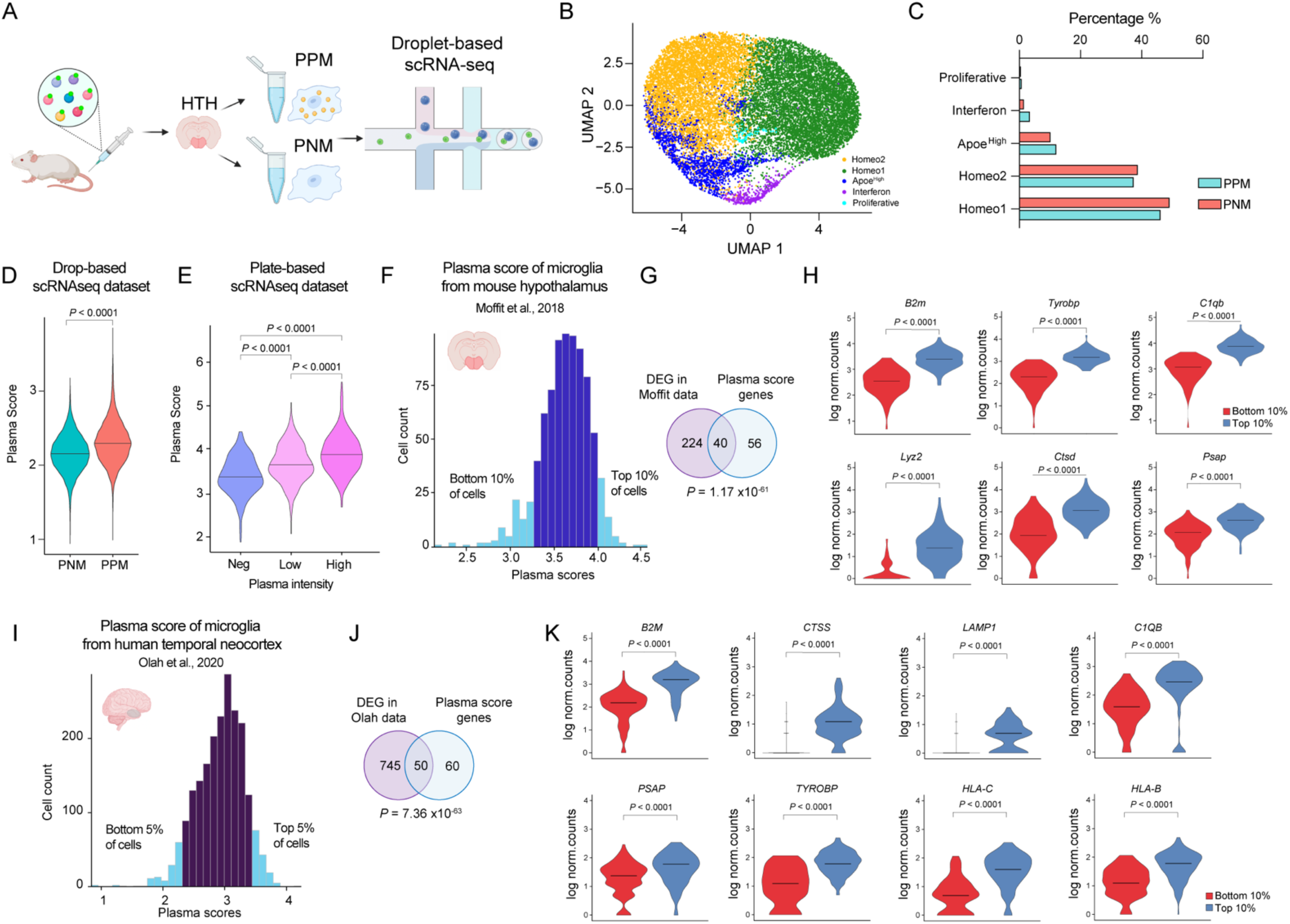
Characterize plasma-positive microglia at single-cell level. A) Experimental scheme for sorting plasma-positive microglia (PPM) and plasma-negative microglia (PNM) from the hypothalamus (HTH) of 3-month-old young mice 20 hours after injecting NHS-Atto647N labeled plasma proteins. The sorted microglia were then used to perform droplet-based single-cell RNA sequencing (scRNA-seq). B) UMAP plot of microglia subtype clusters from the scRNA-seq dataset. Homeo1: Homeostatic 1; Homeo2: Homeostatic 2. C) Normalized percentage of PPM and PNM in each cluster. D) Violin plot of microglia plasma scores for PNM and PPM from the droplet-based mouse scRNA-seq dataset. Data are mean with probability density of plasma score (Wilcoxon Rank-Sum test). E) Violin plot of microglia plasma scores for plasma-negative microglia (Neg), plasma-low microglia (Low) and plasma-high microglia (High) from the index-sorted plate-based mouse scRNA-seq dataset. Data are mean with probability density of plasma score (Kruskal-Wallis test, followed by post-hoc test using use Two-stage linear step-up procedure of Benjamini, Krieger and Yekutieli). F) Histogram showing the distribution of plasma scores for 827 hypothalamic microglia from young mice in a published scRNA-seq dataset^42^. Microglia with the top 10% and bottom 10% of plasma scores are labeled in light blue. G) Differentially expressed genes (DEG) were generated between microglia with the top 10% and bottom 10% plasma scores using the Moffit data^42^. The resulting DEG list was then overlapped with the mouse plasma signature score list (Table S7 and S8; hypergeometric test). The numbers in the Venn diagram indicate the number of genes in the intersections. H) Log-transformed normalized counts of genes were shown for microglia with the top 10% and bottom 10% plasma score from the Moffit data (adjusted *P*-value from Wilcoxon Rank-Sum test and Bonferroni correction). I) Histogram showing the distribution of plasma scores for 2,103 human microglia from temporal neocortex of control group in a published scRNA-seq dataset^43^. Microglia with the top 5% and bottom 5% of plasma scores are labeled in light blue. J) DEGs were generated between microglia with the top 5% and bottom 5% plasma scores using the Olah data^43^. The DEG list was then overlapped with the human plasma signature score list (Table S7 and S8; hypergeometric test). The numbers in the Venn diagram indicate the number of genes in the intersections. K) Log-transformed normalized counts of genes were shown for microglia with the top 5% and bottom 5% plasma score from the Olah data^43^ (adjusted *P*-value from Wilcoxon Rank-Sum test and Bonferroni correction). See also Figure S3-S5, Table S7 and S8.

To further test if the plasma score can distinguish microglia based on relative levels of plasma uptake, we employed a plate-based single-cell transcriptomic profiling approach. Here, we recorded plasma uptake intensity in microglia from dye-labeled plasma-injected mice via flow cytometry and index-sorted them for deep scRNA-seq (Figure S4A). We categorized microglia into three groups: plasma-negative microglia, plasma-low microglia, and plasma-high microglia (Figure S4A-C). Interestingly, plasma score analysis not only showed a significantly higher score in plasma-low microglia than plasma-negative microglia, but also distinguished plasma-high microglia from plasma-low microglia (Figure 3E, Figure S4D). Plate-based data analysis revealed similar DEGs between plasma-positive (including low and high) microglia and plasma-negative microglia, including *Lyz2* and *Apoe* (Figure S4E-G). These data further validate the plasma transcriptional score as a tool to identify uptake of circulatory proteins in microglia.

Next, to ask if the plasma-positive microglia exist in naïve mouse brains, we applied the plasma score to a published dataset where microglia were isolated from naïve mice, without injecting dye-labeled plasma proteins. In the Moffit dataset^42^, where single-nuclei RNA-seq was performed from the hypothalamic preoptic region, some microglia exhibited higher plasma scores, while others displayed lower plasma scores (Figure 3F). The number of differentially expressed genes between the top 10% plasma score-high and bottom 10% plasma score-low microglia largely overlapped with plasma-positive gene signature, including *B2m, Tyrobp, C1qb, Lyz2, Ctsd* and *Psap,* which were upregulated in the top 10% plasma score-high microglia (Figure 3G and H, Figure S5A-C; Table S8), suggesting that the top 10% plasma score-high microglia are very similar to plasma-positive microglia.

Finally, to ask if putative plasma-positive microglia exist in human brains, we applied the plasma score to a published human microglia dataset. The Olah et al. dataset contains microglia isolated from the human temporal neocortex region obtained at autopsy by single-cell RNA sequencing^43^. A comparison between the top 5% plasma score-high cells and bottom 5% plasma score-low cells revealed that genes related to antigen presentation (*B2M, HLA-C* and HLA-B), lysosome function *(CTSS, LAMP1* and *PSAP*) were highly expressed in the top 5% plasma score-high microglia, indicating that these plasma score-high microglia closely resemble plasma-positive microglia (Figure 3I-K, Figure S5D-F; Table S8). Taken together, our meta-analysis suggests a similar microglia population capable of taking up peripheral plasma proteins is also present in human brains.

### Human microglia take up plasma proteins in a brain-region specific manner in chimeric mice model

To further extend our findings to human biology, we sought to assess the capacity of human microglia to take up plasma proteins *in vivo*, using a recently developed CSF1R-G795A human microglia transplanted chimeric mouse model^44,45^ (Figure 4A). EGFP-positive human hematopoietic progenitor cells (HPCs) were bilaterally transplanted into the cortex and dorsal hippocampus region of adult mice (M-CSF^h^ line). 2 weeks later, mice were treated with PLX3397 chow, a potent CSF1R-inhibitor, to deplete and replace endogenous mouse microglia. Subsequently, mice were transitioned to standard chow for 1 month before Atto647N-labled plasma injection. Immunostaining confirmed CNS-wide replacement of murine microglia with EGFP-positive cells (Figure 4B) that exhibit typical microglial morphology and express IBA-1, the human-specific nuclear marker Ku80 (Figure 4C), and a high level of the homeostatic microglial marker P2RY12 (Figure 4D). Twenty hours after Atto647N-labeled plasma injection, human microglia were isolated from micro dissected brain regions for flow analysis. Remarkably, we observed not only uptake of plasma proteins by human microglia but also noted a similar regional heterogeneity in plasma uptake (Figure 4E, Figure S6), as reported in endogenous microglia above (Figure 1C). Specifically, the highest plasma uptake was observed in the hypothalamus (16.9% of total microglia), followed by the thalamus (7.6%) and hippocampus (6.83%), with the lowest uptake in the cerebellum (2.3%). Given that these human HPC-derived microglia were first expanded *in vitro,* then transplanted into a single location in the brain before they migrated and engrafted into distinct regions, we conclude that the specification of microglia into PPM is determined by their local environment through yet unidentified factors.

**Figure 4.**
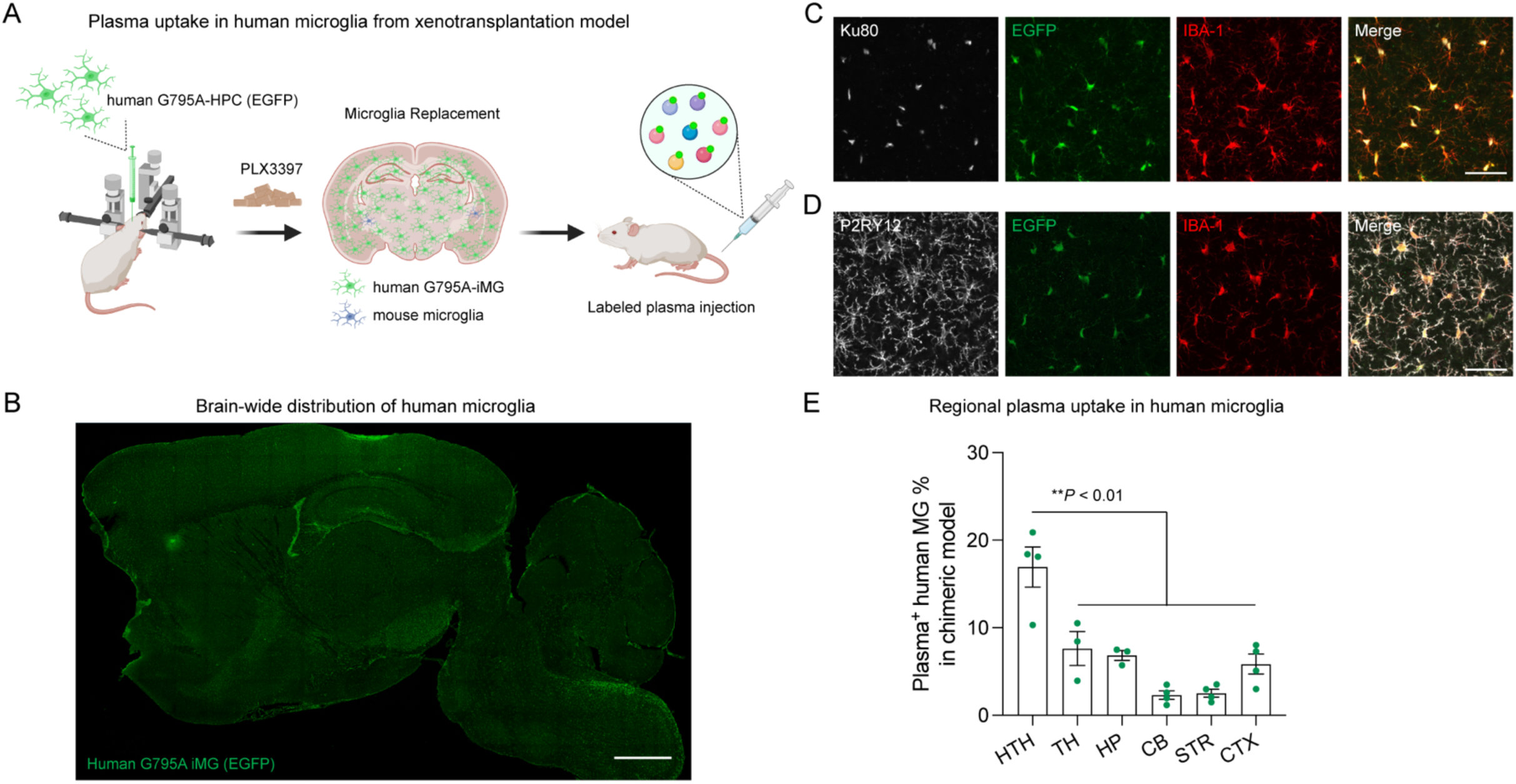
Human microglia take up plasma proteins from blood in a brain-region specific manner in CSF1R-G795A human microglia transplanted chimeric mouse model. A) Experimental scheme of testing plasma uptake in CSF1R-G795A human microglia transplanted chimeric mouse model. The detailed protocol to generate human microglia replacement model was described previously^44,45^. B) Tile image of a sagittal brain section shows the widely distribution of EGFP-positive G795A human microglia in chimeric mice model after microglia replacement. Scale bar, 1000 μm. C) Fluorescence microscopy image shows the transplanted EGFP-positive iMG express microglia marker IBA-1 (red) and human nuclear marker (Ku80, grey). Scale bar, 50 μm. D) Fluorescence microscopy image shows the transplanted EGFP-positive iMG express microglia marker IBA-1 (red) and microglial homeostatic marker (P2RY12, grey). Scale bar, 50 μm. E) Quantification of the percentage of plasma-positive human microglia versus total microglia in six brain regions of adult chimeric mice (n = 3-4 mice per group; mean ± s.e.m.; one-way ANOVA, followed by Tukey’s post hoc test). HPC, Hematopoietic progenitor cells; iPSC-microglia, iMG; HTH, hypothalamus; TH, thalamus; HP, hippocampus; CB, cerebellum; STR, striatum; CTX, cortex. See also Figure S6.

### ApoA-I as a circulatory protein taken up by microglia

The specialized blood-brain barrier (BBB) selectively permits the entry of bloodborne signals into the brain^46^. To identify circulatory proteins taken up by microglia, we developed an unbiased *in vitro* proteomic platform to inform our candidate proteins for *in vivo* assessments. We used NHS-PEG-Biotin to label the entire plasma proteome and filtered the biotin labeled-plasma through an *in vitro* BBB transwell model that contains a single layer of highly confluent mouse endothelial cells^47^, mimicking the *in vivo* transcytosis of proteins across brain endothelial cells. The confluency of the BBB transwell model was confirmed by the low permeability of dextran-FITC (40 kDa and 100 kDa). BBB-filtered plasma proteins collected from the bottom chamber were then added to primary microglia for six hours, before cells were washed, lysed and biotin labeled-plasma proteins taken up by microglia were enriched by streptavidin beads and identified by mass spectrometry (Figure 5A).

**Figure 5.**
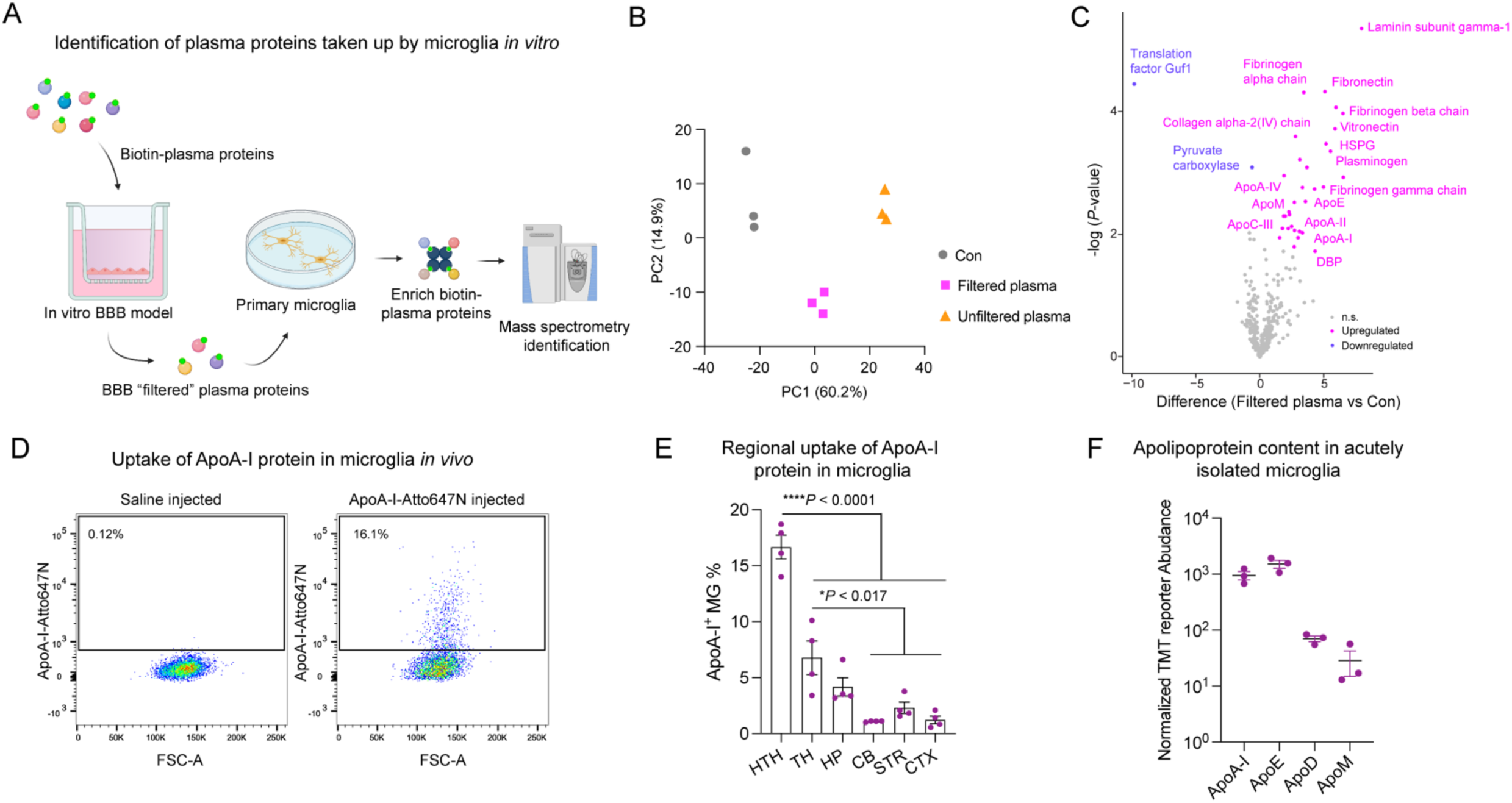
ApoA-I is identified as a circulatory protein that is taken up by microglia *in vivo*. A) Experimental scheme of an unbiased proteomic assay to identify the plasma proteins taken up by microglia *in vitro*. NHS-PEG-Biotin labeled plasma proteins were filtered through an *in vitro* BBB model. The BBB-filtered plasma proteins were then added to primary microglia culture (DIV12) and incubated for six hours. After washing and then lysing the cells, biotin-tagged proteins were enriched using streptavidin beads and analyzed by mass spectrometry using untargeted data-dependent acquisition. B) Principal component analysis (PCA) of proteins enriched from microglia using streptavidin beads after treatment *in vitro* with filtered plasma, unfiltered plasma, and unlabeled plasma (Con) (n = 3 replicates). C) Volcano plot showing proteins enriched in the filtered plasma-treated group (Filtered plasma) compared to the unlabeled plasma-treated group (Con). Proteins downregulated in the filtered plasma-treated group are colored in blue, and proteins upregulated in the filtered plasma-treated group are colored in magenta (*P*-value < 0.05). D) Representative flow cytometry data of the same number of pre-gated microglia (CD45^low^CD11b^+^CD206^-^) from the hypothalamus of NHS-Atto647N labeled ApoA-I protein injected mice compared to those from saline-injected mice. The percentages of ApoA-I positive microglia are shown. E) Quantification of the percentage of ApoA-I-positive microglia versus total microglia in six brain regions of 3-month-old mice (n = 4 mice per group; mean ± s.e.m.; one-way ANOVA, followed by Tukey’s post hoc test). F) Normalized TMT reporter abundance of four apolipoproteins from a published quantitative proteomic dataset of acutely isolated adult mouse microglia (mean ± s.e.m.)^53^. See also Figure S7 and S8, Table S9.

We found that proteins identified from the filtered plasma-treated group are distinct from the unfiltered plasma-treated group (Figure 5B) with 32 proteins enriched and 2 proteins decreased in abundance in filtered plasma group compared with the control group (Figure 5C; Table S9). Many enriched proteins belonged to the Apolipoprotein family or were involved in blood coagulation pathways, including fibrinogen and plasminogen. To determine if these proteins are indeed being taken up by microglia *in vivo*, we individually labeled 10 proteins commercially available in recombinant form with NHS-Atto647N, injected them intravenously into mice, isolated microglia, and looked for fluorescent microglia by flow cytometry (Figure S7). We found Apolipoprotein A-I (ApoA-I, Figure 5D) and Apolipoprotein A-II (ApoA-II, Figure S7G) were robustly taken up by microglia *in vivo*. Additional negative controls were albumin, the most abundant plasma protein (Figure S7C), and transferrin, a protein known to cross the BBB and the basis of brain drug delivery programmes^48^ (Figure S7F) that we have previously shown to be taken up into neurons but not microglia using this method^16^. These results suggest that protein uptake of ApoA-I and ApoA-II into microglia are likely specific. Furthermore, we found that microglia from different brain regions exhibit differential uptake of ApoA-I (Figure 5E), with the highest uptake percentage in the hypothalamus, followed by the thalamus and hippocampus, mirroring the pattern seen with whole plasma proteins (Figure 1C). Our kinetics assay suggests that the highest uptake of ApoA-I occurred twenty hours after intravenous injection (Figure S8A).

ApoA-I is one of the most abundant apolipoproteins in both human and mouse cerebrospinal fluid (CSF)^49,50^ and has previously been reported to enter the central nervous system through the choroid plexus and by crossing the BBB via endocytosis^51,52^. Interestingly, both ApoA-I and ApoA-II are constituents of high-density lipoprotein (HDL) and may share a similar mechanism for entering the brain. As ApoA-I is more abundant than ApoA-II –comprising 70% of the protein carried by HDL, we focused on the role of ApoA-I in microglia as a proof of concept in the current study. ApoA-I is distinct from other apolipoproteins since it is only produced by peripheral organs, primarily the liver and intestine, and it is not expressed by any cells in the brain (Figure S8B and C); in contrast, ApoE and ApoJ both are expressed in peripheral tissues and in the brain. Interestingly, ApoA-I is enriched in published mass spectrometry proteomic data from sorted mouse microglia^53^ (Figure 5F) and can be detected in *ex vivo* primary human microglia^54^, supporting our observations that ApoA-I in microglia originates from peripheral sources. The scavenger receptor BI (SR-BI) which can mediate HDL transcytosis across endothelial cells *in vitro*^55,56^ is also expressed in microglia^29^. Indeed, we found that inhibition of cell surface SR-BI with a blocking antibody against SR-BI significantly reduced the uptake of Atto647N-labeled ApoA-I in primary microglia and brain endothelial cells *in vitro* (Figure S8D and E). These data suggest SR-BI could potentially function in the uptake of ApoA-I into the brain and microglia.

### ApoA-I positively regulates microglial phagocytosis *in vitro* and *in vivo*

To investigate the role of ApoA-I uptake in microglia, we first examined its effect on microglia *in vitro* and found that ApoA-I exhibits an anti-inflammatory effect in lipopolysaccharide (LPS)-induced inflammatory stress by bulk RNA-seq (Figure S9). Administration of ApoA-I inhibits expression of pro-inflammatory genes, including *Cxcl1*, *Ccl3*, and increases expression of the homeostatic genes including *Sall1* in LPS-treated cells. Considering that plasma-positive microglia exhibit higher phagocytic function than plasma-negative microglia, we next tested if ApoA-I also affects microglial phagocytosis *in vitro* using human-induced pluripotent stem cell–derived microglia (iMG)^57^. Using time-lapse imaging of wild-type iMG cultured with 2 µg/ml labeled fibrillar Aβ (fAβ), we observed human iMG gradually take up fAβ over 24-hours (blue) compared to the PBS treated control iMG (green). Surprisingly, adding ApoA-I protein not only enhanced fAβ phagocytosis when added to culture at 0 hours (red), but also alters ongoing microglial phagocytosis when added at the 12-hour point (orange). Notably, microglial accumulation of fAβ following addition of ApoA-I at 0 and 12-hours converge above the blue curve, suggesting ApoA-I is capable of increasing and sustaining human microglial phagocytosis of fAβ (Figure 6A). These results were further validated in BV2 microglia-derived cells which demonstrate enhanced phagocytosis of pHrodo-Zymosan beads and fAβ following ApoA-I treatment (Figure S10A-E). Taken together, these data demonstrate ApoA-I is sufficient to increase the microglial phagocytosis of extracellular material.

**Figure 6.**
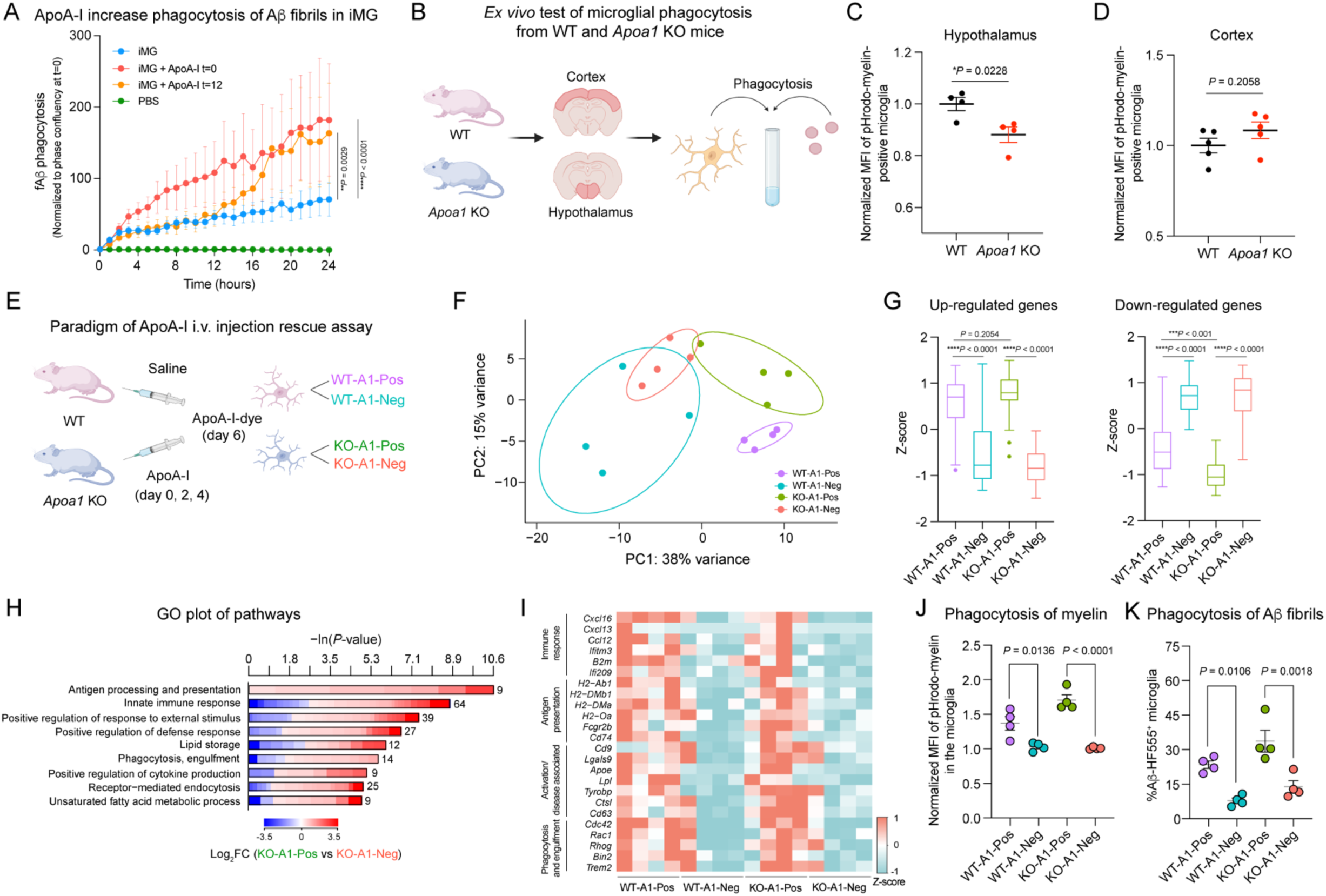
ApoA-I positively regulates microglia phagocytic function both *in vitro* and *in vivo*. A) ApoA-I (100 µg/ml) increase phagocytosis of Aβ fibrils in iMG (ADRC76 WT) when added at t=0 or t=12 hours (n = 6 replicates; two-way ANOVA and Tukey’s post hoc multiple comparisons test; mean ± s.e.m.). B) Experimental design of testing *ex vivo* phagocytic function of microglia, which are isolated from cortex and hypothalamus of adult wild-type (WT) and A*poa1* KO mice. C) Normalized geometric mean fluorescence intensity (MFI) of pHrodo-myelin positive microglia from hypothalamus regions (n= 4 mice per group; unpaired two-tailed *t*-test; mean ± s.e.m.). D) Normalized geometric mean fluorescence intensity (MFI) of pHrodo-myelin positive microglia from cortex regions (n= 5 mice per group; unpaired two-tailed *t*-test; mean ± s.e.m.). E) Experimental design of ApoA-I protein rescue assay. Adult *Apoa1* KO mice were injected 3 times of ApoA-I proteins intravenously on day 0, day 2 and day 4. Wild-type mice of the same age were injected with saline. Both groups were injected with dye labeled ApoA-I (Atto647N-ApoA-I) on day 6. 20 hours later, ApoA-I-Positive and ApoA-I-Negative microglia were isolated from both groups for bulk RNA-seq analysis and *ex vivo* phagocytosis assay. F) PCA plot of bulk transcriptomes of sorted microglia from four groups (n = 4 mice per group). G) Boxplot representation of Z-score transformed expression levels of 50 differentially up- and down-regulated genes when comparing KO-A1-Pos microglia versus KO-A1-Neg microglia (Kruskal-Wallis test, followed by post-hoc test using use Two-stage linear step-up procedure of Benjamini, Krieger and Yekutieli). H) Representative Gene Ontology (GO) enrichment categories of KO-A1-Pos microglia versus KO-A1-Neg microglia. The lengths of the bars represent the negative natural logarithm (ln) of transformed *P*-value obtained using a two-sided Fisher’s exact test. The colors indicate gene-wise log_2_ fold-changes between the two groups. The numbers beside the bars indicate the number of DEGs in that specific GO category. I) Heatmap showing the significant differentially expressed genes between the four groups. The TPM of genes is Z-score transformed (n = 4 mice per group; *P* < 0.05). J) Normalized geometric mean fluorescence intensity (MFI) of *ex vivo* microglial phagocytosis of pHrodo-myelin debris in four groups (n= 4 mice per group; mean ± s.e.m.; One-way ANOVA, followed by Tukey’s post hoc test). K) Percentage of *ex vivo* microglial phagocytosis of Aβ-fibrils (Aβ-HF555) in four groups (n= 4 mice per group; One-way ANOVA, followed by Tukey’s post hoc test). See also Figure S9 and S10, Table S10 and S11.

To examine whether ApoA-I is also a necessary regulator of microglial phagocytosis *in vivo*, we utilized *Apoa1* global knock out mice (*Apoa1* KO), in which microglia no longer have access to circulatory ApoA-I. We isolated microglia from the cortex and the hypothalamus regions of 9-month-old *Apoa1* KO and wild-type mice and tested them for *ex vivo* phagocytosis (Figure 6B). Interestingly, we observed a significant decrease in microglial phagocytosis of pHrodo-myelin in the hypothalamus of *Apoa1* KO mice (Figure 6C), while there was no difference in microglia from the cortex (Figure 6D). These data are consistent with the observation that microglia from the hypothalamus preferentially take up plasma proteins including ApoA-I compared with cortical microglia in wild-type mice (Figure 5E), indicating that ApoA-I can regulate microglia in specific brain regions including phagocytosis in hypothalamic microglia.

Lastly, to determine if circulatory ApoA-I is sufficient to restore microglial function in *Apoa1* KO mice to a more wild-type-like state, we intravenously injected ApoA-I protein three times, two days apart, into adult *Apoa1* KO mice (Figure 6E). One day before taking down the mice, we injected dye-labeled ApoA-I into both wild-type mice and *Apoa1* KO mice. This enabled us to isolate and compare ApoA-I-positive and ApoA-I-negative microglia from hypothalamus region of the same mouse and to analyze their transcriptome using bulk RNA-seq and assess their phagocytic function *ex vivo*. The ApoA-I protein-positive cells in the *Apoa1* KO mice effectively took up ApoA-I resulting from our injection. The PCA plot and Z-score analysis of the top 50 differentially expressed genes (Figure 6F and G) indicated that the transcriptome of ApoA-I-positive microglia from *Apoa1* KO mice (KO-A1-Pos) is similar to ApoA-I-positive microglia from wildtype mice (WT-A1-Pos), as opposed to ApoA-I-negative microglia from *Apoa1* KO mice (KO-A1-Neg). Expression of genes associated with pathways, including antigen presentation (*H2-DMa, H2-DMb1*), innate immune response (*Ccl16, Ccl12*), and phagocytosis (*Cdc42, Trem2*), are rescued in ApoA-I-positive microglia from *Apoa1* KO mice (KO-A1-Pos) (Figure 6H, 6I and Figure S10F; Table S10). Consistent with the transcriptomic analysis, when these microglia were incubated with pH-sensitive fluorescent myelin (phrodo-myelin) or Aβ fibrils *ex vivo*, ApoA-I-positive microglia from *Apoa1* KO mice (KO-A1-Pos) exhibited a significantly higher phagocytic capacity than ApoA-I-negative microglia from *Apoa1* KO mice (KO-A1-Neg) (Figure 6J and K), suggesting ApoA-I uptake in these microglia boosts their engulfment and phagocytosis at the functional level. In summary, intravenous injection of ApoA-I can effectively revert the transcriptional state of *Apoa-1* KO microglia toward a wild-type phenotype and enhance their phagocytic functions.

## Discussion

The circulatory system connects every organ in our body delivering not only essential oxygen and nutrients but a plethora of regulatory molecules. In spite of the BBB, the brain seems to be similarly sensitive to circulatory factors with plasma from physically exercised mice^8,58^ or from young mice^11^ increasing brain function while factors from aged mice induce inflammation and aspects of aging in young brains^5,59^. Furthermore, extravasation of plasma proteins, due to a compromised BBB, notably fibrinogen^60,61^, can trigger neuroinflammation and possibly contribute to neurodegenerative and neurological diseases^62,63^. Most of these general effects are not understood at the cellular and molecular level but given that circulatory proteins can enter the brain readily under physiological conditions and be taken up into different neural cell types^16^, it is possible that circulatory proteins modulate function of these cell types and thus regulate higher brain function. Here, we identify a stable population of microglia specialized to take up circulatory proteins in a region-specific manner under physiological conditions. We comprehensively analyze this state at transcriptomic, proteomic, metabolomic, and lipidomic levels. Convergence across these different biological measures, together with functional assays, reveals that plasma-positive microglia exhibit higher metabolic activity and enhanced phagocytic capacity, appearing specialized for antigen presentation. We also identify ApoA-I as a key plasma protein taken up by microglia to regulate microglia phagocytic functions.

Advances in single-cell genomics technologies have profoundly enhanced our understanding of microglial heterogeneity, revealing distinct microglial states across various physical and disease contexts in both rodent models and human brains. Most classifications have been based on gene expression signatures^28,29,32,64–69^. Our plasma-positive microglia (PPM) signature, defined here from microglia in the hypothalamus of young healthy brains — a relatively less characterized region — differs from DAMs/MGnD defined in Alzheimer’s disease or other neurodegenerative diseases, despite some overlapping genes such as *Apoe*, *Axl*, *Tyrobp* and *Lyz2*. Unique PPM signature genes that differ from DAMs/MGnD include *Ccl5, Ctss, Psap, C1qb and Ifitm3* (Table S11). Using multi-omics profiling coupled with functional assays, PPM are characterized by innate immune response, chemotaxis, and antigen processing and presentation. They display higher metabolic activity and phagocytic activity than PNM. Given recent findings on the impact of microglia-T cell crosstalk on aging^70^ and neurodegenerative disease progression, especially in Alzheimer’s disease (AD)^71,72^, plasma-positive microglia may actively sense signals in the blood, engaging in the uptake of proteins in a healthy state, and responding more rapidly and robustly to stress to maintain brain homeostasis. Moreover, the increased metabolic activity and enhanced phagocytic function of PPM may be beneficial in scenarios such as amyloid-beta plaque clearance in AD^73^ and myelin debris removal in aging and demyelinating diseases^68,74^. Exploring the role of PPM in aging and various disease contexts will likely provide new insights into these hypotheses in the future.

The uptake of plasma proteins by microglia decreases notably in the hypothalamus and hippocampus during aging. Additionally, our previous work showed that the composition of plasma proteins undergoes substantial changes cross the entire human lifespan^75^. Given the important roles of the hypothalamus in regulating essential physical functions including sleep, body temperature, blood pressure, and mood^76,77^, as well as the pivotal role of the hippocampus in learning and memory, the potential impact of age- and disease-related alterations in protein uptake by microglia on neural circuit function is intriguing. Understanding the interactions between PPM and neighboring cells in these regions could shed light on how these alterations further affect neural circuit function.

Since similar physical distances between PPM or PNM and the BBB were observed, other factors apart from the proximity to the vasculature may contribute to the brain regional heterogeneity in microglial uptake of plasma proteins. The presence of permeable fenestrated capillaries in the hypothalamus^78^ may enable the brain to sense and respond to sensory and secretory signals in the blood. Microglia in the hypothalamus might also be exposed to a higher concentration of plasma proteins compared to other regions, leading to a higher uptake. However, considering that only a specific subset of plasma proteins can enter the brain, especially in other regions such as thalamus and hippocampus with a tight BBB, protein first need to traverse endothelial cells and then be taken up by microglia. This mechanism likely depends on the expression of receptors on both cell types, thereby also influencing the regional variation in microglial uptake of plasma proteins. Additionally, there are emerging concepts of plasma protein uptake at the blood-CSF barriers and subsequent biodistribution along perivascular pathways into the brain from the CSF^79^.

Principally, the functional heterogeneity of microglia and the abundance of PPM across different brain regions may be regulated by intrinsic programming, such as epigenetic regulation of their clearance activity^30^ or by extracellular cues from the local environment. For instance, microglia differentially suppress neuronal activation in the grey matter and white matter of the striatum in response to IL34 or CSF1 from neighboring cells^80^. Our findings with transplanted human microglia suggest that the state and specialization to become plasma-positive microglia is not cell intrinsic and can be induced by their spatial distribution within the brain and potentially by exposure to circulatory proteins. Notably, the consistency in brain region heterogeneity of plasma uptake between mouse and human microglia underlines the broader relevance of our findings to microglial biology.

The role of ApoA-I in high-density lipoprotein (HDL) metabolism is well studied in the periphery and has been shown to provide protection from atherosclerotic cardiovascular disease by facilitating reverse cholesterol transport, reducing inflammation, and promoting endothelial nitric oxide (NO) synthase activity^81^. However, the functional impact of ApoA-I on the brain is less clear. Our work indicates that ApoA-I treatment reduces inflammation in microglia under lipopolysaccharide (LPS), suggesting a common anti-inflammatory role of ApoA-I in multiple cell types. Our work also shows that ApoA-I increases microglial phagocytosis of Aβ fibrils and myelin debris. Previous studies found that ApoA-I can form a complex with Aβ fibrils and prevent Aβ-induced neurotoxicity^82^ and HDL reduces soluble brain Aβ levels in symptomatic APP/PS1 mice^83^. Interestingly, enhancing cholesterol efflux by ApoA-I treatment restores monocyte-derived tumor-associated macrophage phagocytosis, thereby boosting anti-tumor immunity in glioblastoma^84^. Additionally, genetic *Apoa1* overexpression reduces neuroinflammation, mitigates cerebral amyloid angiopathy, and improves cognitive function in mouse models of AD while genetic deletion of *Apoa1* has the opposite effects^85–87^. Given that microglia and macrophages are closely related cell types it is conceivable that ApoA-I mediates its functional benefits on microglia by modulating cholesterol metabolism. Future studies should also explore possible relationships between lipid droplet formation in microglia^88^, amyloidosis, and ApoA-I.

In the current study we highlighted the uptake of ApoA-I into PPM, but it is highly likely that many other circulatory proteins enter these specialized cells and some of them may regulate their function. Advances in protein labeling and enrichment tools should help accelerate the discovery of such proteins and set the stage to explore their possible roles in microglial function. Enhancing the entrance of beneficial plasma proteins, or inhibiting the entrance of detrimental proteins, may offer new opportunities to regulate the communication between periphery and the CNS and lead to a new class of CNS therapies. Furthermore, harnessing and engineering endogenous brain-entering plasma proteins into so-called “brain shuttles” may facilitate the delivery of therapeutic cargos to the brain.

### Limitations of the study

Our study assesses plasma uptake by microglia mostly through chemical tagging of plasma proteins, which may modify protein surface charge, activity, binding, and trafficking, and thus may influence their uptake. Another limitation is that we cannot conclusively determine at this point whether the molecular signature of PPM is governed by local cues from their environment, the uptake of circulatory proteins, or both. Lastly, ApoA-I is typically bound to lipid particles, and it is very likely the dye-labeled ApoA-I injected intravenously will bind to lipids and HDL. Similar to challenges studying ApoE, it is thus currently not feasible to separate the effects of lipid-free from lipid-bound ApoA-I *in vivo*.

## Acknowledgments

We are grateful to M. Schoof, T. Iram, A. Tsai, H. Oh, R. Palovics, N. Khoury, other past and current members of the Wyss-Coray lab for valuable discussions, D. Channappa, H. Zhang, K. Dickey for laboratory management, M. Block from Indiana University School of Medicine for providing the BV2 cell line, G. Randolph from Washington University School of Medicine, St. Louis for reagents and suggestions, Y. Gao from Westlake University for helping with Imaris data processing, B. Carter and W. Wang from the Stanford Veterans Affairs FACS core for flow cytometry technical expertise, H. Shin for help with droplet-based scRNA-seq sample loading step, K. Kim for technical assistance with bioinformatics data. We thank A. Atebi, A. Kolodkin, and P. Shi for critically reading the manuscript. In addition, the authors express their gratitude and respect to all animals sacrificed in this study. Schematics were created with Biorender.com.

We acknowledge the following funding support: To T.W.-C.: Simons Foundation, Hong Kong Center for Neurodegenerative Disease, D. H. Chen Foundation, and NIH (R01AG072255); To N.L.: AHA-Allen Brain Health and Cognitive Impairment Cross-Network Collaborative Grants (23BHCICG1188316); To JP.C.: NIH (T32 AG073088); To J.H.: NIH (T32 AG00096); To J.L.: the US Department of Defense (HT9425-23-1-0879) and NIH (AG059694); To T.H.C.: the Innovation and Technology Commission (ITCPD/17-9); To A.C.Y.: NIH Director’s Early Independence Award (1DP5OD033381) and the Burroughs Wellcome Fund Career Awards at the Scientific Interface; To Q.L.: NIH (R01AG078512); To M.A.-R.: NCL Foundation (NCLStiftung). M.A.-R. is a Stanford Terman Fellow and a Pew-Stewart Scholar for Cancer Research, supported by the Pew Charitable Trusts and the Alexander and Margaret Stewart Trust; To M.B.-J.: NIH (U19 AG06970101) and NIH (AG016573). N.L. and I.G. are MAC3 Dementia and Ageing Fellows supported by MAC3 Impact Philanthropies. S.M.S. was supported by a Stanford Bio-X Fellowship.

## Author contributions

N.L. and T.W.-C. conceptualized the study. N.L. designed and performed most of the experiments, except for those indicated below. P.M.-L., O.H., A.S., E.T., and N.L. analyzed the bulk RNA-seq data. A.S., E.T., O.H., and N.L. analyzed the scRNA-seq data. JP.C., J.H., H.D., and N.L. conducted the iMG and human MG transplanted mice experiments under the supervision of M.B.-J.. N.L., A.S., E.T., J.S., I.G., L.L., and J.L. performed microglia isolation and downstream characterizations. W.D. and N.L. conducted metabolic and lipidomic profiling and data analysis under the supervision of M.A.-R.. W.Z., N.L., N.-S.T., and P.S.W. conducted TimsTOF profiling and data analysis under the supervision of T.H.C.. N.L., S.M.S, and S.R.S performed *in vitro* proteomic assay. A.Y.C. and Q.L. provided critical intellectual contributions. N.L. wrote the manuscript. N.L. and T.W.-C. edited the manuscript with input from all authors. T.W.-C. supervised the study. All authors read and approved the final manuscript.

---

**Figure S1.**
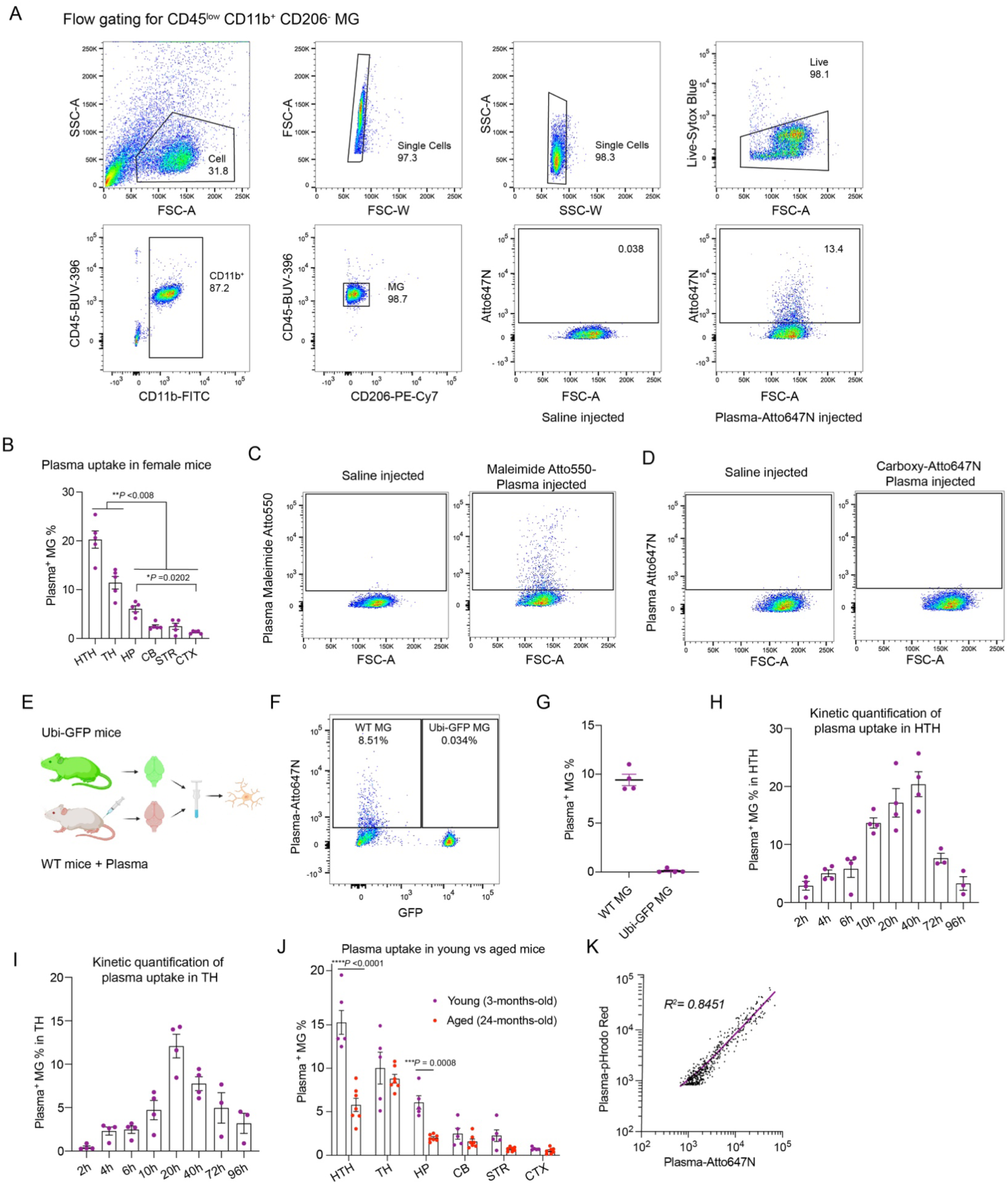
Detailed characterizations of plasma uptake in microglia. A) Representative flow cytometry plots showing the gating strategy for live microglia (CD45^low^CD11b^+^CD206^-^) used in this study. B) Quantification of the percentage of plasma-positive microglia versus total microglia in six brain regions of 3-month-old female mice (n = 5 mice per group; mean ± s.e.m.; one-way ANOVA, followed by Tukey’s post hoc test). C) Representative flow cytometry plots showing post-gated hypothalamic microglia (CD45^low^CD11b^+^CD206^-^) from Maleimide Atto550 labeled plasma-injected mouse compared to saline-injected mouse. Unlike NHS-Atto647N, which labels proteins via lysine residues, Maleimide Atto550 dye label proteins via thiol residues. A decent amount of plasma positive microglia was observed using Maleimide Atto550 dye. D) Representative flow cytometry plots showing pre-gated hypothalamic microglia (CD45^low^CD11b^+^CD206^-^) from carboxy-Atto647N labeled plasma injected mouse compared to saline-injected mouse. When plasma proteins are incubated with a fluorophore without a covalent conjugative moiety (carboxy-Atto647N), no plasma-positive microglia were observed. E) Experimental scheme. Hypothalamus tissue from Ubi-GFP mice uninjected was mixed with hypothalamus tissue from wild-type mice 20 hours after injecting plasma-Atto647N. Mixed tissues were minced, dounce dissociated and analyzed by flow cytometry. F) Representative flow cytometry plots showing post-gated hypothalamic microglia (CD45^low^CD11b^+^CD206^-^) from mixed tissues. Plasma-positive microglia only exist in GFP negative cells (from wild-type mice) but do not exist in GFP-positive cells (from Ubi-GFP mice). G) Quantification of the percentage of plasma-positive microglia from wild-type mice and Ubi-GFP mice (n = 4 mice per group; mean ± s.e.m.). H and I) Kinetic quantification of the percentage of plasma-positive microglia in the hypothalamus (H, HTH) and thalamus (I, TH) at different time points after injecting the same dosage of plasma-Atto647N into young mice (n= 3-4 mice per group; mean ± s.e.m.). J) After injecting dye labeled plasma proteins, quantification was performed 20 hours later for the percentage of plasma-positive microglia versus total microglia in six brain regions of young mice (3-month-old) and aged mice (24-month-old) (n = 5 mice for young group and n = 7 mice for aged group; mean ± s.e.m.; two-way ANOVA, followed by Sidak’s post hoc test). K) Correlation analysis of Atto647N-plasma positive microglia versus pHrodo Red-plasma positive microglia of the mice that received twice plasma injections (derived from flow cytometry data in Fig. 1L). HTH, hypothalamus; TH, thalamus; HP, hippocampus; CB, cerebellum; STR, striatum; CTX, cortex; ns, not significant.

**Figure S2.**
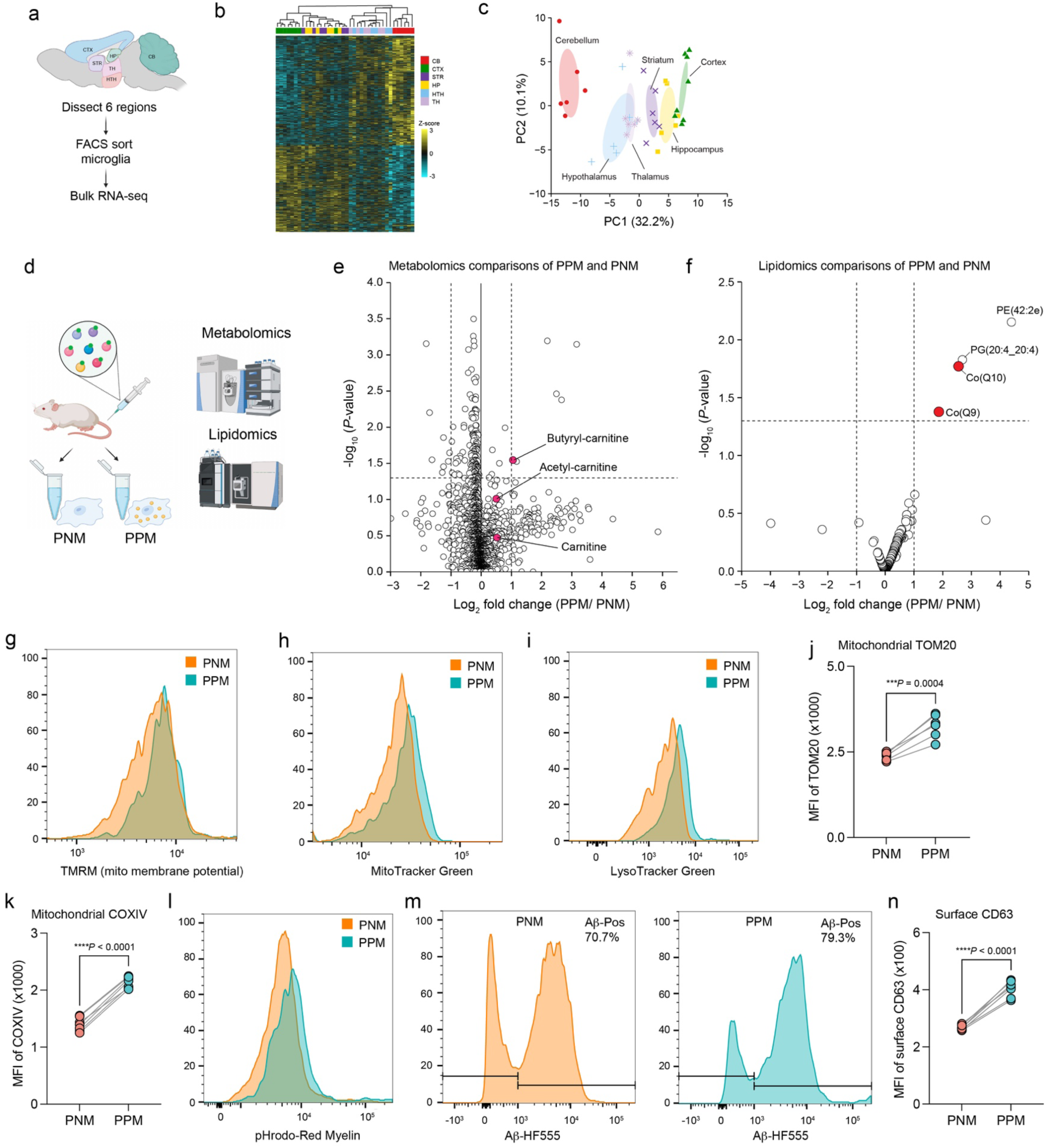
Microglia bulk RNA-seq from six brain regions; metabolic, lipidomic and functional characterizations of plasma-positive microglia. A) Experimental scheme for FACS-sorting microglia from six major brain regions of young mice (3-month-old). The sorted microglia were then used for bulk RNA-seq. B) Heatmap showing 1521 candidate genes that differentiate microglia by their brain region of origin. Column labels indicate brain region. Genes that passed the statistical cutoff (*P* < 0.05; DESeq2’s Two-sided Wald test) in at least one comparison between any pair of regions was included. C) Principal component analysis (PCA) of microglia from six different brain regions (n = 8 mice for CTX; n = 6 mice for CB, STR, HP, TH and HTH). D) Experimental scheme for sorting plasma-positive microglia (PPM) and plasma-negative microglia (PNM) from the hypothalamus of 3-month-old young mice 20 hours after injecting NHS-Atto647N labeled plasma proteins. The sorted microglia were then used to perform metabolomic and lipidomic assays. 50,000 microglia were sorted per replicate from pooled mice. E) Volcano plot comparing the metabolomics data from sorted plasma-positive microglia (PPM) and plasma-negative microglia (PNM) shows that butyryl-carnitine increase significantly in PPM, along with a trend towards an increase in two other carnitine related metabolites, acetyl-carnitine and carnitine. Horizontal line indicates a *P*-value of 0.05, and vertical dotted lines indicate a fold change of 2. Each dot represents a metabolite (n = 5 biological replicates, unpaired two-sided *t*-test). Data are in Table S5. F) Volcano plot comparing the lipidomic data from sorted plasma-positive microglia (PPM) and plasma-negative microglia (PNM) shows that Co(Q10) and Co(Q9) increase significantly in PPM. Horizontal line indicates a *P*-value of 0.05, and vertical dotted lines indicate a fold change of 2. Each dot represents a lipid (n = 6 biological replicates, unpaired two-sided *t*-test). Data were acquired in negative ion mode and normalized to internal lipid standards for the best-matched lipid class. Data are in Table S6. G-I) Representative flow histograms of TMRM (mitochondria membrane potential, G), MitoTracker Green (H), and LysoTracker Green (I) staining in plasma-positive microglia (PPM) and plasma-negative microglia (PNM) isolated from hypothalamus of mice, 20 hours after injecting them with NHS-Atto647N labeled plasma proteins. J and K) Geometric mean fluorescence intensity (MFI) of intracellular TOM20 staining (J) and intracellular COXIV staining (K) in PPM and PNM isolated from hypothalamus of mice, 20 hours after injecting them with NHS-Atto647N labeled plasma proteins (n =6 mice per group for TOM20; n =7 mice per group for COXIV; paired two-sided *t*-test). L) Representative flow histograms of *ex vivo* phagocytosis of pHrodo-Red labeled myelin in PPM and PNM isolated from hypothalamus of mice, 20 hours after injecting them with NHS-Atto647N labeled plasma proteins. M) Representative flow histograms of *ex vivo* phagocytosis of HF555 labeled Aβ in PPM and PNM isolated from hypothalamus of mice, 20 hours after injecting them with NHS-Atto647N labeled plasma proteins. N) Geometric mean fluorescence intensity (MFI) of surface CD63 staining in PPM and PNM isolated from hypothalamus of mice, 20 hours after injecting them with NHS-Atto647N labeled plasma proteins (n =7 mice per group; paired two-sided *t*-test). HTH, hypothalamus; TH, thalamus; HP, hippocampus; CB, cerebellum; STR, striatum; CTX, cortex.

**Figure S3.**
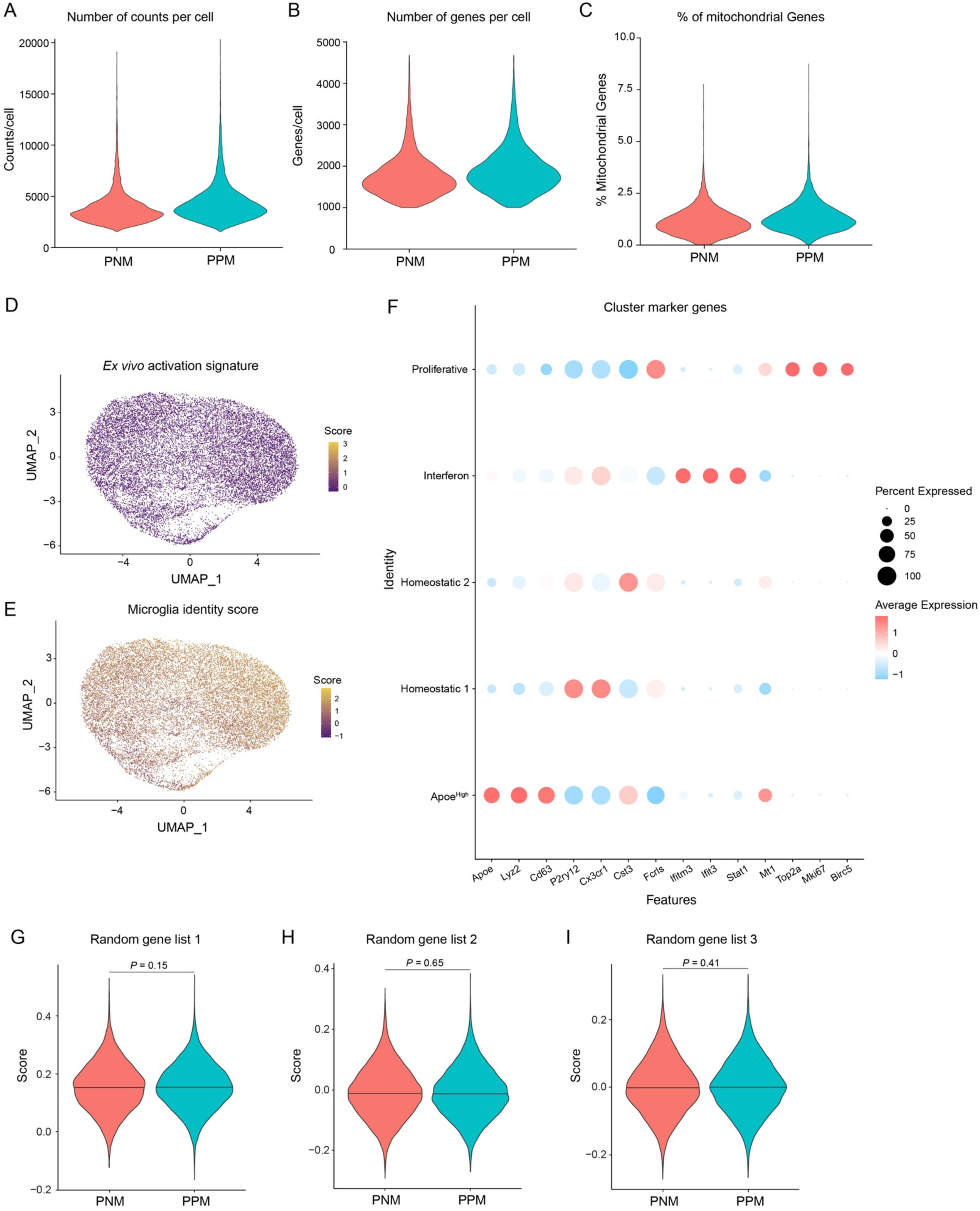
Quality control metrics for droplet-based scRNA-seq dataset. A-C) Violin plots of quality control (QC) metrics for droplet-based scRNA-seq datasets. The data is split by plasma-positive microglia (PPM) and plasma-negative microglia (PNM) and are presented as a distribution of QC metrics (count per cell (A), genes per cell (B), percentage of mitochondrial genes(C)). D) Expression feature plot for the *ex vivo* activation signature in the dataset. *Ex vivo* activation scores are defined as a function of gene expression for core *ex vivo* activation genes reported previously^39^ (Table S7). E) Expression feature plot for microglia identity score in the dataset. Microglia identity scores are defined as a function of gene expression for core microglia genes identified previously^39^ (Table S7). F) Dots plot for cluster marker genes. The size of each circle represents the percentage of total microglia expressing the gene. The transformed average expression levels are shown using different colors. G-I) Randomly select 3 different lists of 100 genes from the microglial bulk RNA-seq dataset in Figure 2A and use these genes to calculate a score for each cell. Violin plot of random gene scores for PNM and PPM from the droplet-based mouse scRNA-seq dataset. Data are mean with probability (Wilcoxon Rank-Sum test).

**Figure S4.**
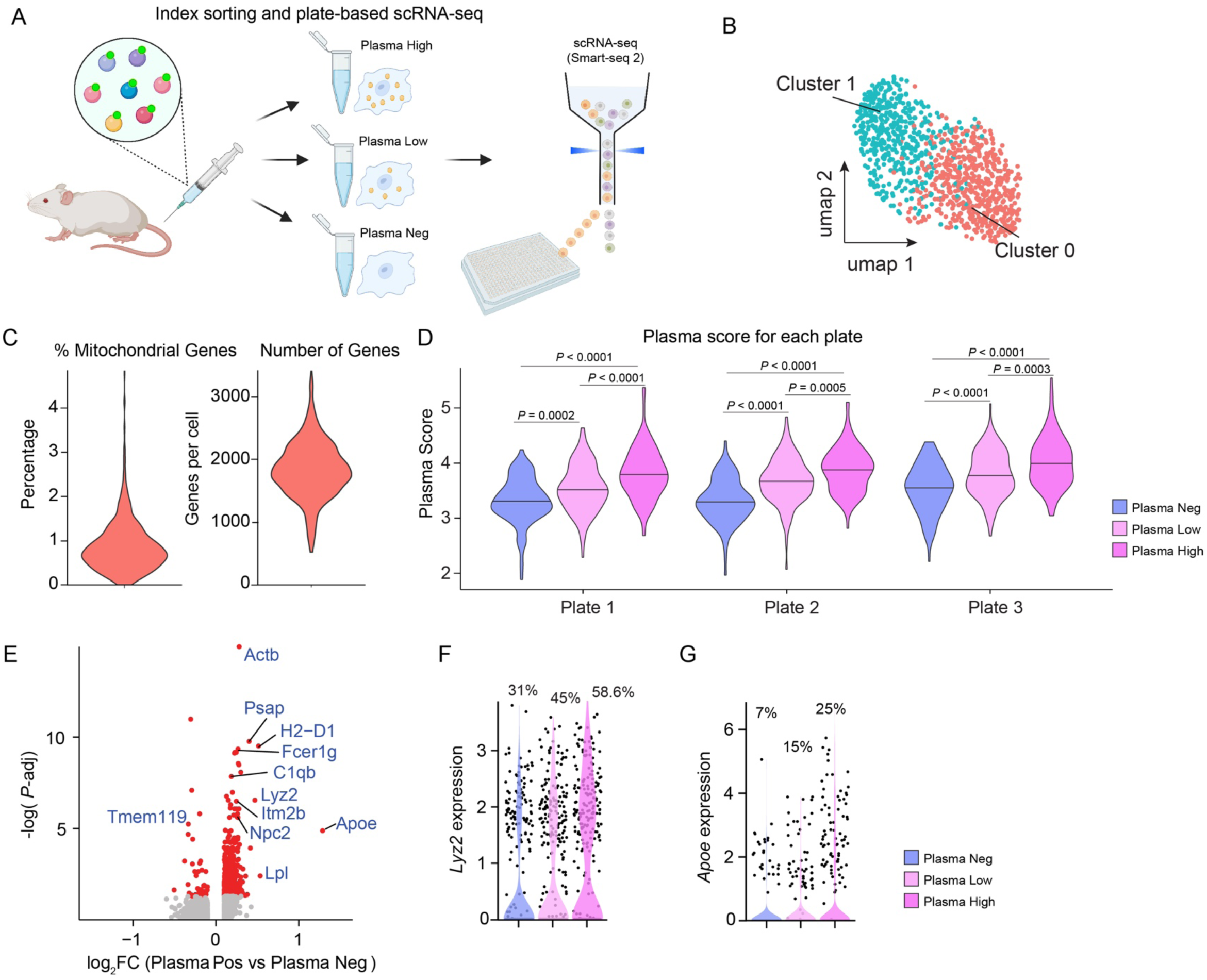
Characterizations of plasma high- and plasma-low microglia using plate-based scRNA-seq. A) Experimental scheme for indexed sorting PPM and PNM into 384 well plates. Microglia are isolated from 3-month-old young mice, 20 hours after injecting NHS-Atto647N labeled plasma proteins. Based on the intensity of plasma protein within each cell, microglia were assigned into plasma negative (plasma neg), plasma low and plasma high groups. The sorted microglia were then used to perform plate-based single-cell RNA sequencing (scRNA-seq). B) UMAP plot of microglia clusters from the scRNA-seq dataset. C) Violin plots of the percentage of mitochondria genes in each cell and the number of nfeature (genes) detected in each cell. D) Violin plot of microglia plasma scores for plasma-negative microglia (Plasma Neg), plasma-low microglia (Plasma Low) and plasma-high microglia (Plasma High) from the index-sorted plate-based mouse scRNA-seq dataset. The cells are split into 3 plates. Data are mean with probability density of plasma score (Kruskal-Wallis test, followed by post-hoc test using use Two-stage linear step-up procedure of Benjamini, Krieger and Yekutieli). E) Volcano plot of differentially expressed genes (DEGs) between plasma-positive microglia (including both plasma high and plasma low) and plasma-negative microglia. Significantly DEGs are labeled with red color (*P-*adjust < 0.05, using Benjamini–Hochberg FDR correction). F and G) Violin plots of normalized expression level of *Lyz2* (F) and *Apoe* (G) in each group. The percentage values indicate the percentage of cells in each group expressing the gene.

**Figure S5.**
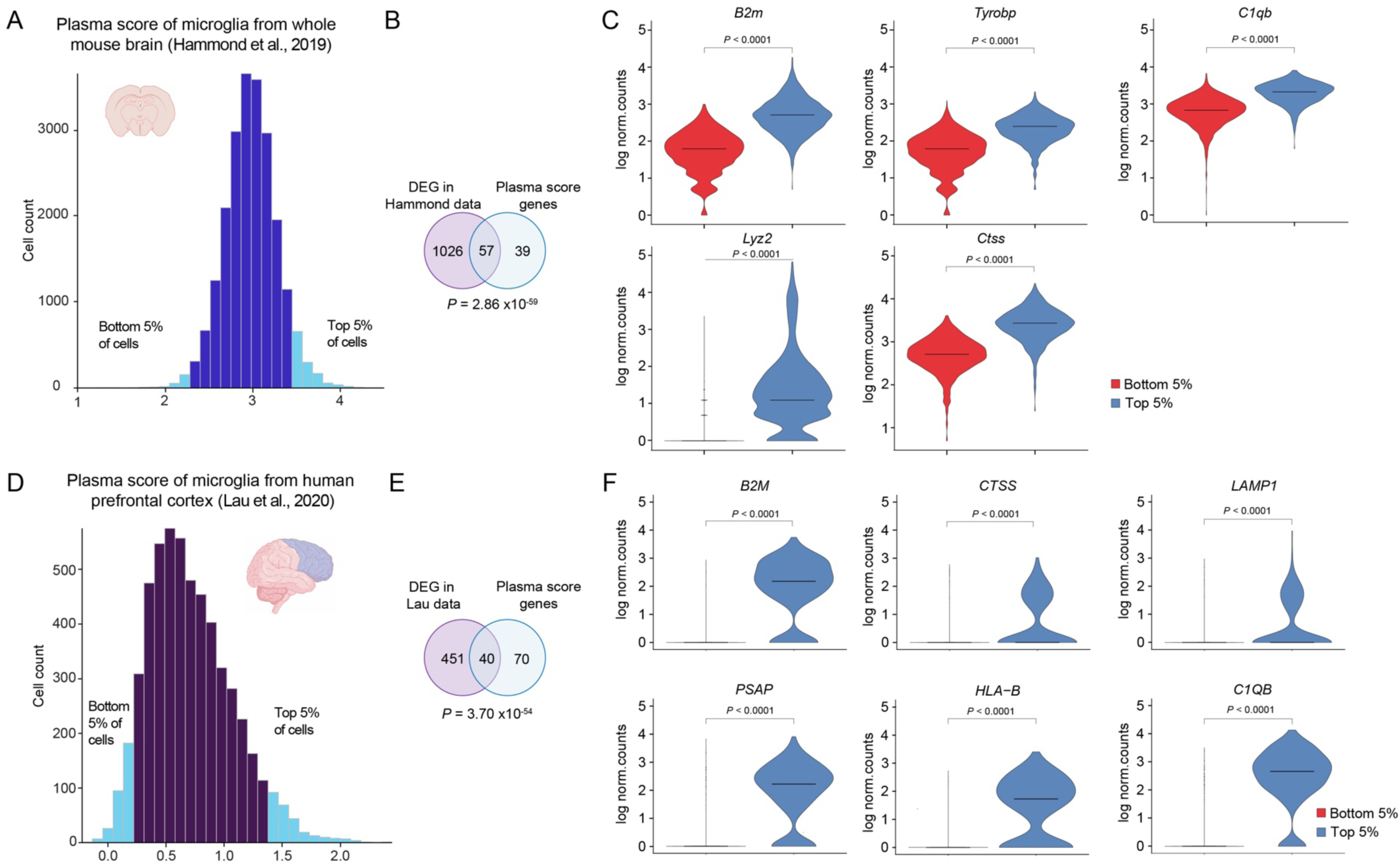
Plasma score analysis in additional published scRNA-seq datasets. A) Histogram showing the distribution of plasma scores for 16,782 microglia from the whole brain of young mice (postnatal 100) in a published scRNA-seq dataset^29^. Microglia with the top 5% and bottom 5% of plasma scores are labeled in light blue. B) Differentially expressed genes (DEG) were generated between microglia with the top 5% and bottom 5% plasma scores using the Hammond data^29^. The resulting DEG list was then overlapped with the mouse plasma score list (Table S7 and S8; hypergeometric test). The numbers in the Venn diagram indicate the number of genes in the intersections. C) Log-transformed normalized counts of genes were shown for microglia with the top 5% and bottom 5% plasma score from the Hammond data^29^ (adjusted *P*-value from Wilcoxon Rank-Sum test and Bonferroni correction). D) Histogram showing the distribution of plasma scores for 2,733 human microglia from prefrontal cortex of the control group in a published single-nucleus RNA sequencing dataset^89^. Microglia with the top 5% and bottom 5% of plasma scores are labeled in light blue. E) DEGs were generated between microglia with the top 5% and bottom 5% plasma scores using the Lau data^89^. The DEG list was then overlapped with the human plasma score list (Table S7 and S8; hypergeometric test). The numbers in the Venn diagram indicate the number of genes in the intersections. F) Log-transformed normalized counts of genes were shown for microglia with the top 5% and bottom 5% plasma score from the Lau data^89^ (adjusted *P*-value from Wilcoxon Rank-Sum test and Bonferroni correction).

**Figure S6.**
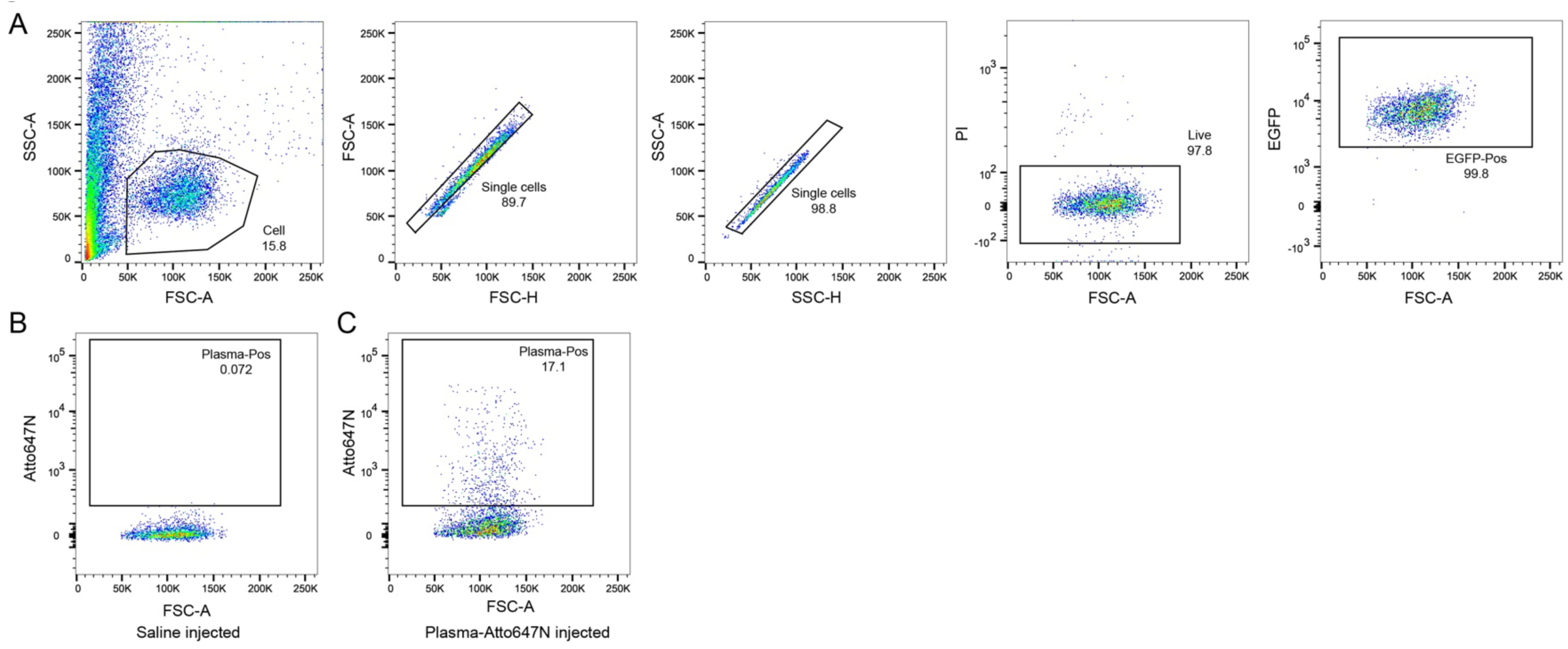
Flowcytometry data for human microglia gating in CSF1R-G795A human microglia transplanted chimeric mouse model. A) Representative flow cytometry plots showing the gating strategy for live and EGFP-positive human microglia from transplanted mice. B and C) Representative flow cytometry data of pre-gated human microglia from saline-injected mouse (B) and from Atto647N labeled plasma-injected mouse (C). The percentages of plasma-positive microglia are shown.

**Figure S7.**
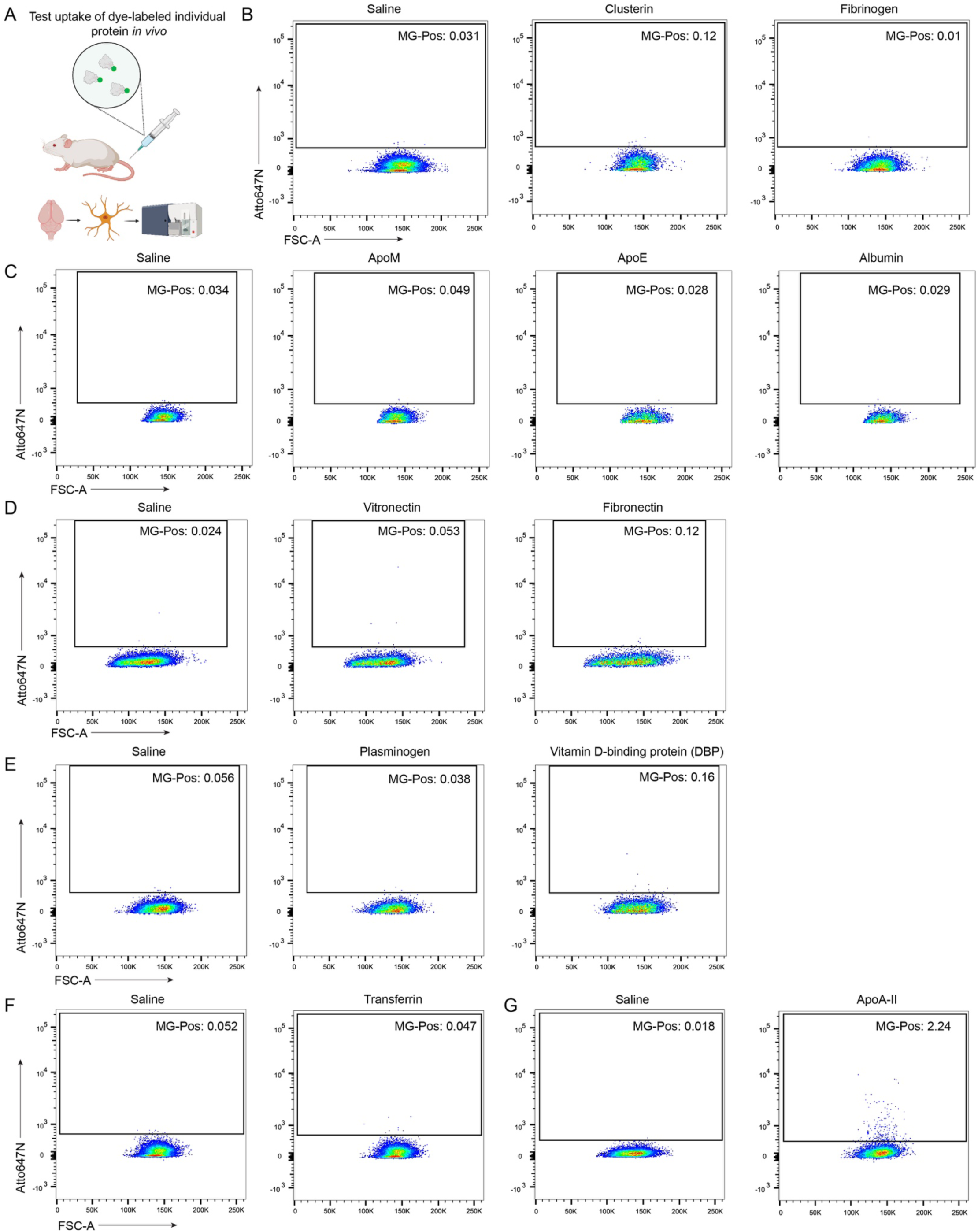
Test the uptake of candidate proteins in microglia *in vivo* by single protein Atto647N labeling and injection. Representative flow cytometry data of pre-gated microglia (CD45^low^CD11b^+^CD206^-^) in the hypothalamus region of NHS-Atto647N labeled individual protein-injected mouse comparing to a saline-injected mouse (3-month-old). The percentages of protein-positive microglia are shown. A) Experimental scheme for testing the uptake of NHS-Atto647N labeled individual protein in microglia by flow cytometry. B) Mice injected with Clusterin or Fibrinogen are compared to the saline-injected group. C) Mice injected with labeled ApoM, ApoE or Albumin are compared to the saline-injected group. D) Mice injected with Vitronectin or Fibronectin are compared to the saline-injected group. E) Mice injected with Plasminogen or Vitamin D-binding protein (DBP) are compared to the saline-injected group. F) Mouse injected with Transferrin is compared to the saline-injected group. G) Mouse injected with ApoA-II is compared to the saline-injected group.

**Figure S8.**
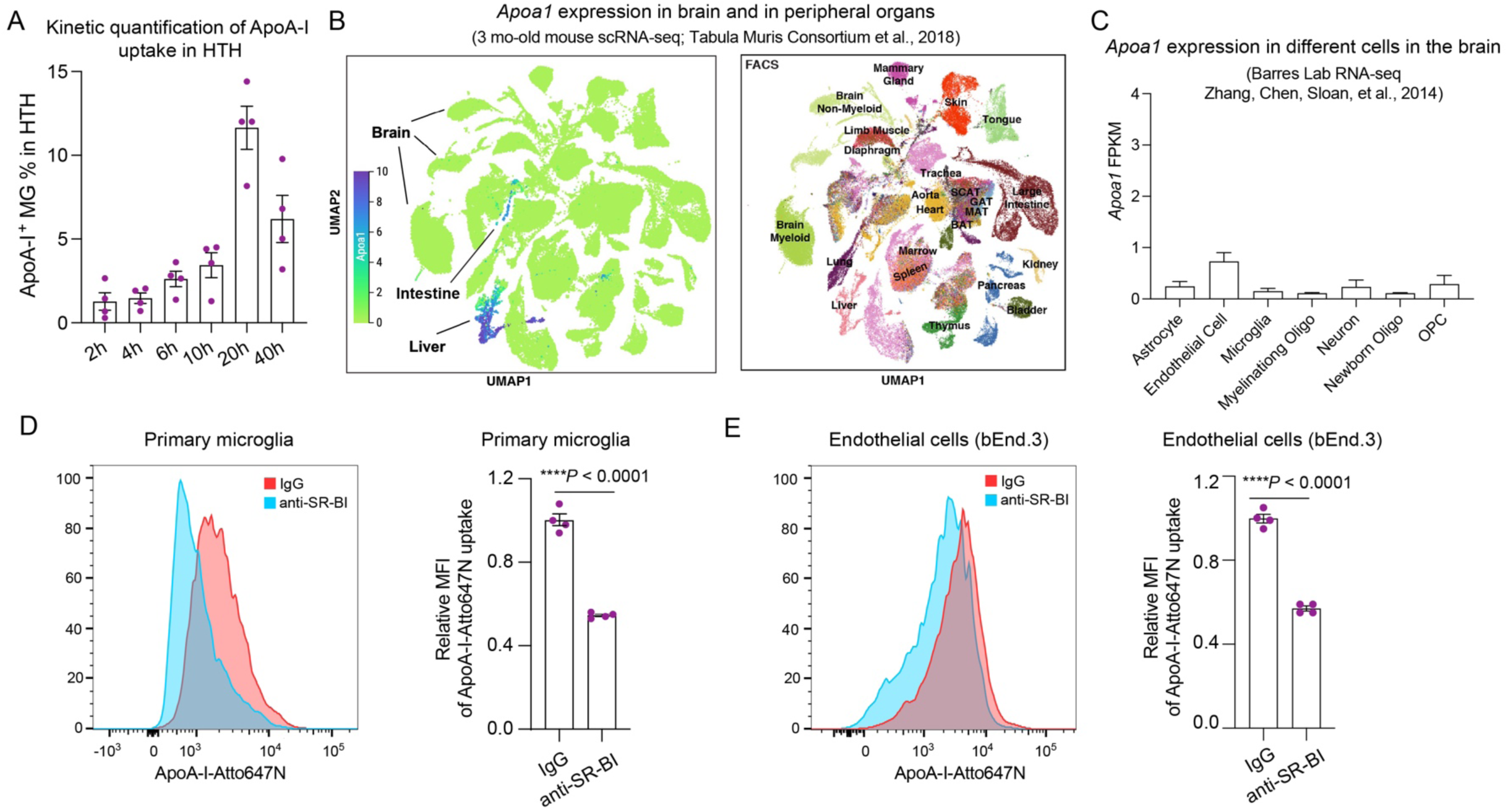
Expression of *Apoa1* in published mouse RNA-seq datasets and SR-BI antibody blockades ApoA-I uptake *in vitro*. A) Kinetic quantification of the percentage of ApoA-I-positive microglia in the hypothalamus at different time points after injecting the same dosage of ApoA-I-647N into young mice (n=4 mice per group; mean ± s.e.m.). B) UMAP plot showing scRNA-seq analysis of *Apoa1* expression in cells from 20 different mouse tissues. *Apoa1* is expressed in the liver and intestine but is absent from brain cells (both myeloid and non-myeloid). Data from the Tabula Muris Consortium^90^. C) *Apoa1* expression in various cell types of the young mouse brain (bulk RNA-seq), showing that *Apoa1* is barely expressed in any brain cells. Data from Barres laboratory bulk RNA-seq^91^ (http://www.brainrnaseq.org/). D) Flow histograms and relative mean fluorescence intensity (MFI) of ApoA-I-Atto647N uptake with a blocking antibody against SR-BI or with IgG control in primary microglia (n = 4; unpaired two-sided *t*-test; mean ± s.e.m.). E) Flow histograms and MFI of ApoA-I-Atto647N uptake with a blocking antibody against SR-BI or with IgG control in brain endothelia cells (bEnd.3) (n = 4; unpaired two-sided *t*-test; mean ± s.e.m.).

**Figure S9.**
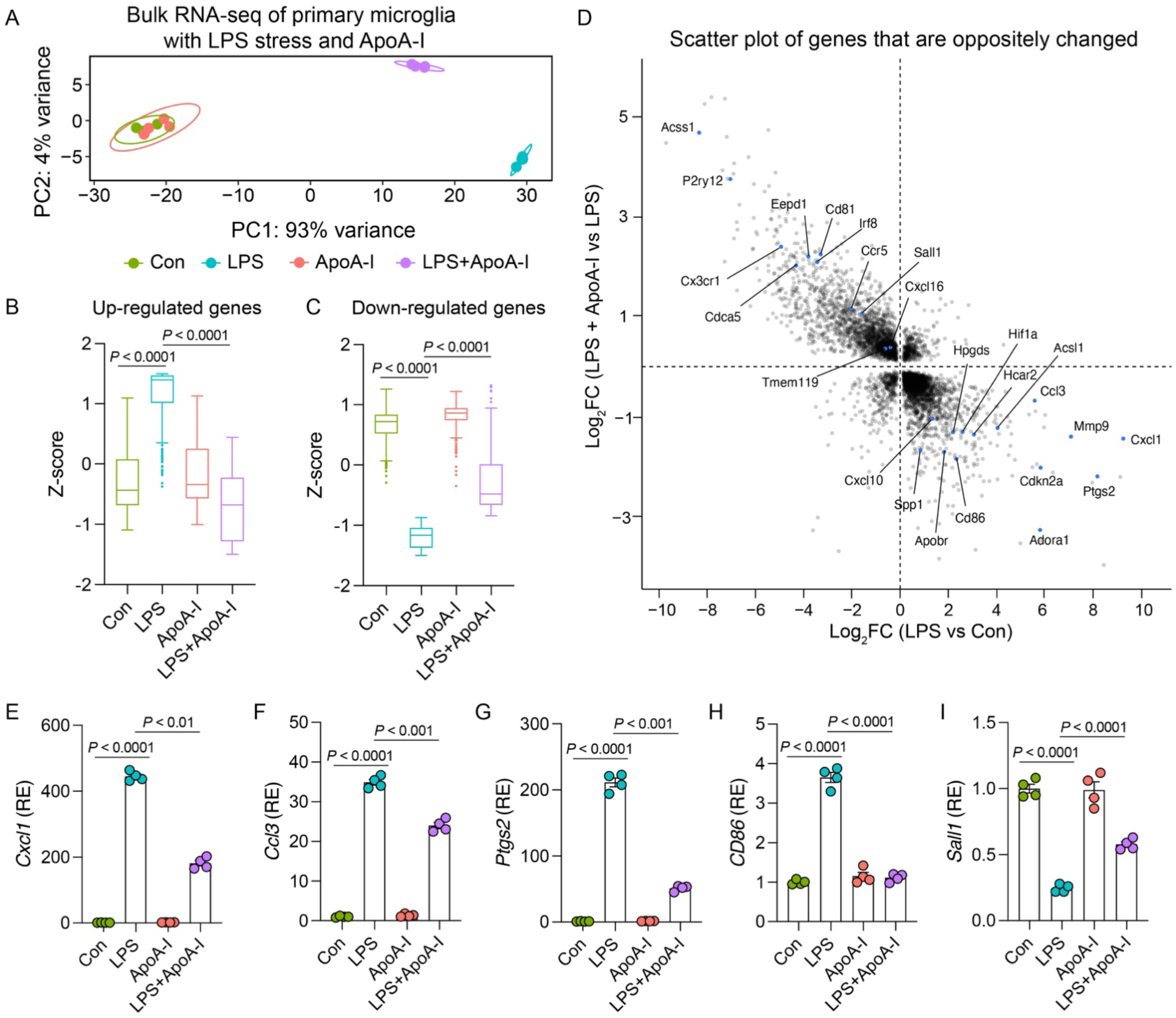
ApoA-I exhibits anti-inflammatory effect in lipopolysaccharide (LPS)-induced inflammatory stress *in vitro*. A) Principal component analysis (PCA) plot of bulk transcriptomes of primary cultured microglia treated with LPS (100 ng/ml) and/or ApoA-I proteins (100 μg/ml) for 20 hours (n = 4 replicates per group). B and C) Boxplot representation of Z-score transformed expression levels of the top 300 differentially up-(B) and down-regulated genes (C) when comparing the LPS-treated group to the LPS+ApoA-I-group (Kruskal-Wallis test, followed by post-hoc test using use Two-stage linear step-up procedure of Benjamini, Krieger and Yekutieli). D) Scatter plot displaying significantly differentially expressed genes in the LPS versus Con comparison (X-axis) and the LPS+ApoA-I versus LPS comparison (Y-axis). Genes located in the upper-left and bottom-right quadrants exhibit a reversed expression pattern upon ApoA-I treatment. E-I) Relative expression (RE) of transcripts per kilobase million (TPM) of representative genes from the bulk RNA-seq dataset (n = 4 replicates per group; mean ± s.e.m.; *P* values are determined by DESeq2).

**Figure S10.**
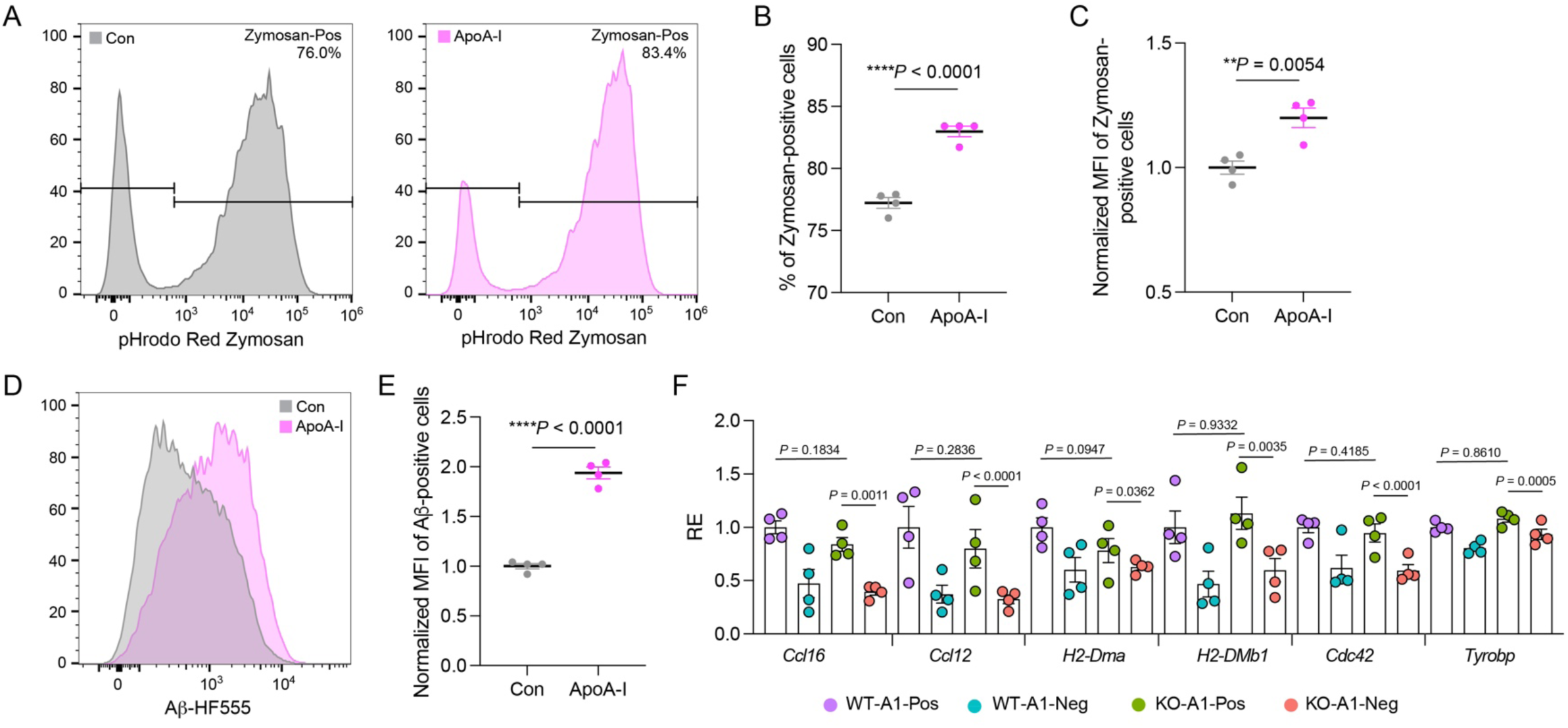
ApoA-I increase phagocytosis of Zymosan beads and Aβ fibril by BV2 cells. A) Representative flow histograms of phagocytosis of pHrodo-red Zymosan beads by BV2 cells with or without ApoA-I treatment (100 μg/ml). B) Percentage of Zymosan-positive cells in each group (n=4 replicates; unpaired two-sided *t*-test; mean ± s.e.m.). C) Normalized geometric mean fluorescence intensity (MFI) of Zymosan-positive cells in each group (n=4 replicates; unpaired two-sided *t*-test; mean ± s.e.m.). D) Representative flow histograms of phagocytosis of Aβ fibrils (HF555-labeled Aβ) by BV2 cells with or without ApoA-I treatment (100 μg/ml). E) Normalized geometric MFI of Aβ-positive cells in each group (n=4 replicates; unpaired two-sided *t*-test; mean ± s.e.m.). F) Relative expression (RE) of transcripts per kilobase million (TPM) of representative genes from bulk RNA-seq dataset of ApoA-I rescue assay (n = 4 mice per group; mean ± s.e.m.; *P* values are determined by DESeq2).

## Method

### Animals

Young C57BL/6J male and female mice (3-months-old) were purchased from the Jackson Laboratories (stock #000664). Aged C57BL/6J male mice (24-months-old) were obtained from the National Institute on Aging rodent colony. Ubi-GFP mice (C57BL/6-Tg(UBC-GFP)30Scha/J, stock #004353) and *Apoa1* KO mice (B6.129P2-Apoa1tm1Unc/J, #002055) were ordered from the Jackson laboratory. M-CSF^h^ mouse line was purchased from Jackson Laboratories (stock #017708) and contains *Rag2* and *Il2rg* deletions and humanized M-CSF^h^. Mice were housed under a 12-hour light-dark cycle and provided water and standard chow *ad libitum*. All animal procedures were approved by the VA Palo Alto Committee on Animal Research, the institutional administrative panel of laboratory animal care at Stanford University and the University of California, Irvine Institutional Animal Care and Use Committee. All experiments used male mice unless otherwise indicated.

### Plasma collection and labeling

Blood was collected by terminal transcardiac bleeding of wild-type C57BL/6J mice (3-month-old, unless otherwise indicated), using 250 mM EDTA (Thermo Fisher Scientific) as an anticoagulant. EDTA plasma was isolated by blood centrifugation at 1,000 g for 15 min at 4 °C, no brake. Plasma was pooled from 40 mice per batch, aliquoted, and stored at −80 °C.

Plasma protein molarity was approximated to be 750 μM. Amine-reactive N-hydroxysuccinimide (NHS) ester moieties, thiol/sulfhydryl-reactive Maleimide moieties or amine-reactive STP ester moieties were used for labeling proteins. The labelling ratios were determined empirically for each specific label: for fluorescence dye, NHS-Atto 647N, NHS-Atto 550, Carboxy-Atto 647N, and Maleimide Atto 550 (ATTO-TEC) was added at 0.8 −1.3 x molar ratio. pHrodo iFL Red STP Ester (Thermo Fisher Scientific, P36010) was added at 3.3x molar ratio and plasma labeling was done following the manufacturer’s instructions. For *in vitro* labelling plasma proteome and mass spectrometry assay, Biotin-PEG4-NHS Ester (Click Chemistry Tools) was added at 15× molar ratio.

NHS dyes were incubated with plasma for 1.5 h rotating at room temperature (RT) and Biotin-PEG4-NHS Ester was incubated over night at 4 °C before PBS dialysis overnight. The next day, 50 mM Tris pH 8.0 was added for 10 min at RT for quenching, and the labelled plasma was washed extensively with Amicon Ultra-15 Centrifugal Filter Unit (Millipore Sigma, 10 kDa cutoff), and was subsequently washed four times with Zeba Spin Desalting Columns, 7K MWCO cutoff (Thermo Fisher Scientific) for clean-up. Plasma was further concentrated with Amicon Ultra-0.5 Centrifugal Filter Unit (Millipore Sigma, 3 kDa cutoff) for retro-orbital injection. Plasma concentrations were measured using the Nanodrop spectrophotometer (Thermo Fisher Scientific) and were stored at 4 °C.

Unless otherwise indicated, 20 mg of labeled plasma proteins were injected retro-orbitally at a volume of 150 µl into 3-month-old recipient mice (0.67 mg/g body weight, in line with previous studies using HRP)^92,93^. Control mice were injected with the same volume of saline. Unless otherwise indicated, transcardiac perfusion with 1x PBS was performed twenty hours later, followed by tissue dissection method as indicated below to isolate microglia. For measuring plasma uptake in aged mice, plasma from 24-month-old mice was used for labeling. For measuring plasma uptake in female mice, plasma from 3-month-old female mice was used for labeling.

### Individual protein labeling and *in vivo* microglia uptake assay

ApoA-I protein (Athens Research &Technology, 16-16-120101-LEL), ApoA-II (Athens Research & Technology, 16-16-120102), ApoM (LSbio, LS-G14868), ApoE (Sigma, SRP4760), Albumin (Sigma, 126674), Clusterin (Sino Biological, 50485), Fibrinogen (Abcam, ab92791), Vitronectin (Sino Biological, 50585-M08H), Fibronectin (Corning, 354008), Plasminogen (Innovative research, IMSPLG1MG), Vitamin D-binding protein (DBP) (Athens Research &Technology, 16-16-070307), and Transferrin (Sigma, T0665) were used for labeling.

Individual protein was incubated with NHS-Atto647N dye for 1.5 h rotating at RT at 1.2x molar ratio. 50 mM Tris pH 8.0 was added for 10 min at RT to quench the reaction. Labeled protein was cleaned once with Zeba Spin Desalting Columns, 7K MWCO cutoff (Thermo Fisher Scientific, 89882), and was concentrated with Amicon Ultra-0.5 Centrifugal Filter Unit (Millipore Sigma, 3 kDa cutoff, UFC500324) for retro-orbital injection. 1 μg labeled protein was loaded on a protein gel and check the successful labeling by in-gel Westerns (LiCOR Odyssey). 0.5-1 mg individual labeled protein was injected into a 3-months-old mouse and twenty hours later, hypothalamic microglia was dissociated and analyzed by flow assay. For the kinetic assay of ApoA-I uptake *in vivo*, 1 mg of labeled ApoA-I protein was injected into individual 3-months-old mouse (40 mg/kg), and the percentage of ApoA-I uptake in microglia was assessed at different time points after injection.

### Immunohistochemistry and quantitative analysis

Mice brain tissue processing and immunofluorescence experiments were performed as described previously^16,32^. Brains were isolated and post-fixed in 4% (w/v) PFA overnight at 4 °C before preservation in 30% (w/v) sucrose in PBS. Brains were sectioned at a thickness of 40 μm, and sections were stored in cryoprotective medium at −20 °C. Free-floating sections were washed three times in PBS followed by 1h blocking in 10% donkey serum and 0.3% Triton X-100, before incubation at 4 °C with the following primary antibodies: CD68 (Bio-Rad, MCA1957, 1:500); Iba1 (Abcam, ab5076, 1:500); Iba1 (Fujifilm Wako, 019-19741, 1:500); CD102 (ICAM2, Bio-Rad, 553326, 1:100); Ku80 (abcam, ab130752, 1:200); GFP (Cell signaling technologies, 2955s, 1:500); P2RY12 (Sigma, HPA014518, 1: 300).

After primary antibody incubation, sections were washed three times in PBS and incubated with the Alexa Fluor-conjugated secondary antibodies (1:500, Invitrogen) for 2 hours at RT. Sections were washed once in PBS and incubated with Hoechst 33342 (1:2,000; Thermo Fisher Scientific) for 5 min and mounted on microscope slides with ProLong Gold (Thermo Fisher Scientific, P36934) before imaging on a confocal laser scanning microscope (Zeiss LSM880). Age-related autofluorescence was quenched with TrueBlack Plus lipofuscin Autofluorescence quencher (Biotium, 23014).

ImageJ (National Institutes of Health) was used for quantitative analysis. To quantify the distance of plasma-positive microglia and plasma-negative microglia to the nearest vessel, Tiled image of hypothalamus regions from 4 brain slices per mouse were captured. Maximum projection of the z-stack across the whole hypothalamus regions was analyzed with ImageJ to manually calculate the distance of microglia to the nearest vessels. 3D reconstruction of confocal image stacks (acquired at ×63 magnification) of plasma-positive microglia were converted to 3D images with the surface-rendering feature of Imaris BitPlane software (9.9).

### Microglia isolation

For most experiments unless otherwise indicated, microglia were isolated by an ice-cold dounce homogenize method as previously described^28,32^. Mice were perfused with 1x PBS and six brain regions including hypothalamus, thalamus, hippocampus, striatum, cortex, and cerebellum were dissected. Brain tissues were chopped and homogenized using a dounce homogenizer in 2 ml of cold medium A (HBSS, 15 mM HEPES, 0.5% glucose and 1:500 DNase I), filtered through a 70-µm cell strainer, rinsed with 6 ml medium A and centrifuged at 400 × g for 5 min. Myelin removal beads and columns from MACS separation system (Miltenyi, beads #130-096-433, columns #130-042-401) were used to remove myelin by incubating 50 µl beads with cells in 450 µl MACS buffer (1x PBS, 2 mM EDTA and 0.5% BSA). After myelin removal, cells were resuspended in FACS buffer (1x PBS, 2 mM EDTA, 1% FBS and 25 mM HEPES). The samples were stained with FC (CD16/CD32, BD, 553141, 1/100) for 5 min on ice and then CD11b-FITC (Biolegend, 101206, 1/100), CD45-BUV395 (BD, 564279, 1/100) and CD206-PE-Cy7 (Biolegend, 141720, 1/100) for 30 min on ice, followed by adding 3 ml FACS buffer and centrifuged at 400 × g for 5 min. Dead cells were excluded by staining with Sytox Blue dead cell stain (1:1,000, Invitrogen, S34857) in the final FACS buffer. Cells were analyzed using an Aria 3.1 (BD Biosciences) with FACSDiva (BD Biosciences).

For measuring mitochondria and lysosome, after antibody staining and spinning down, cell suspension was incubated with TMRM dye (mitochondrial membrane potential indicator, Thermo Fisher Scientific, I34361, 100 nM), MitoTracker Green (Thermo Fisher Scientific, M7514, 1/1000), or LysoTracker Green DND-26 (Thermo Fisher Scientific, L7526, 75 nM) for 15-30 min at 37 °C in the dark. After washing and resuspension, cells were analyzed by flow cytometry.

For surface staining of CD63, after FC blocking for 5 min, CD63-PE (Thermo Fisher Scientific, 12-0631-82, 1/100) was added together with antibodies against CD11b and CD45. For intracellular staining of TOM20 and COXIV, after surface antibodies (CD11b and CD45) incubation for 30 min, Leucoperm kit (Bio-Rad, BUF09) was used for cell fixation and permeabilization. TOM20-CoraLite-488 (Thermo Fisher Scientific, CL48811802100UL), CoXIV-CoraLite-488 (Thermo Fisher Scientific, CL48811242100UL) were added with permeabilization medium. After washing and resuspension, cells were analyzed by flow cytometry.

For the middle-aged *Apoa1* KO mice rescue experiment, we used an enzymatic method to isolate microglia to obtain higher cell yield^68,94^. Transcription inhibitor Actinomycin D (Sigma, A1410, 45 μM) was added to the dissociation solution and enzyme mix to prevent *ex vivo* transcription. In brief, brain tissues were dissociated with the Neural Tissue Dissociation Kit (Papain, Miltenyi, 130-092-628) and filtered through a 70-μm cell strainer (Corning). Debris removal was done using debris removal solution (Miltenyi, 130-109-398). RNase inhibitor (Takara, 2313B, 2 µl/ml) was added to the buffer. After myelin removal, cells were resuspended in FACS buffer. The samples were stained with FC (CD16/CD32, BD, 553141, 1/100) for 5 min on ice and then CD11b-FITC (Biolegend, 101206, 1/100), CD45-BUV395 (BD, 564279, 1/100) and CD206-PE-Cy7 (Biolegend, 141720, 1/100) for 30 min on ice, followed by adding 3 ml FACS buffer and centrifuged at 400 × g for 5 min. Dead cells were excluded by staining with Sytox Blue dead cell stain (1:1,000, Invitrogen) in the final FACS buffer. Cells were sorted using an Aria 3.1 (BD Biosciences) with FACSDiva (BD Biosciences).

### Microglia *ex vivo* phagocytosis assay

Microglia *ex vivo* phagocytosis assay was performed as previously reported with some modifications^74,95^. Myelin debris was isolated from homogenized mice brain using a published sucrose density gradient ultracentrifugation method^96^. To label myelin with pHrodo iFL Red STP Ester (Thermo Fisher Scientific, P36010), 2.5 uL pHrodo-Red SE (6.67 mg/µl), 11 uL of 1 M sodium bicarbonate were added to 100 µg myelin in 100 ul volume and incubate for 60 min at RT. After 3 times wash with PBS, the pHrodo-Red labeled myelin was resuspended in 100 ul PBS and stored at −80°C for phagocytosis assay. HiLyte Fluor 555-labeled Aβ_1-42_ peptide (Aβ-HF555) was obtained from AnaSpec (AS-60479-01) and was reconstituted as suggested by manufacturer with 1.0% ammonium hydroxide, followed by PBS (pH=7.4) to 100 μM and aggregated at 37°C for 5 days. Following the protocol to dissociate microglia, after myelin removal step, cells were spun down and the cell suspension stained with FC blocking for 5 min, then the antibodies against microglia (CD11b and CD45) in FACS buffer on ice for 30 min. After washing, cells were mixed with pHrodo-Red labeled myelin (50 μg/ml in DPBS + 0.5% BSA) or Aβ-HF555 (200 nM in DPBS + 0.5% BSA) and incubated at 37 °C for 60 min with gentle mixing. Cell suspensions were washed and analyzed on a flow cytometer (Aria 3.1, BD Biosciences) with FACSDiva (BD Biosciences). The intensity of pHrodo-myelin^+^ microglia and the percentage of Aβ-HF555^+^ microglia were analyzed.

### BV2 phagocytosis assay

Mouse microglial BV2 cell line was a gift from Dr. Michelle Block and was originally reported from F. Bistoni^97^. Cells were maintained in DMEM supplemented with 10% FBS and antibiotics (Pen/strep, Gibco, 15140122), 10 mM glutamax (Gibco) under standard culture conditions (95% relative humidity with 5% CO_2_ at 37 °C). Adherent cells were split using 1× TrypLE (Gibco).

For phagocytosis assay, cells were plated at 60 k cells/well in 24 well plate. ApoA-I protein (Athens Research &Technology, 16-16-120101-LEL, 100 µg/mL) or vehicle solution was added to the culture medium for 20 hours. After that, phrodo Red Zymosan beads (P35364, Thermo Fisher Scientific, 30 μg/ml) or Aβ-HF555 (200 nM) were added to the medium for 2 hours and 1 hour, separately. Following cargo incubation, cells were washed with 1x PBS, stained with Sytox blue dead cell stain, and analyzed by FACS (MA900 sorters, Sony).

### Bulk RNA-seq library preparation

For sorting plasma-positive microglia and plasma-negative microglia from young mice and performing bulk RNAseq, microglia were isolated by ice-cold dounce homogenize method as described above. RNase inhibitor (Takara, 2313B, 2 µl/ml) was added to the disassociation buffer. 1,000 microglia were sorted into sorted into RLT lysis buffer (Qiagen) with 1% 2-mercaptoethanol (Sigma, M6250) and frozen at −80 °C. For sorting microglia from different brain regions from young mice and performing bulk RNA-seq, microglia were isolated by ice-cold dounce homogenize method as described above. 6,000 microglia were sorted into RLT lysis buffer (Qiagen) with 1% 2-mercaptoethanol (Sigma, M6250) and frozen at −80 °C.

Total RNA was isolated using a RNeasy Plus Micro kit (Qiagen, 74034). cDNA and library syntheses were performed in house using the Smart-seq2 protocol as previously described with the following modifications^8,37^: Extracted RNA was reverse-transcribed using SmartScribe (Takara, 639538) and TSO (Exiqon, 5’-AAGCAGTGGTATCAACGCAGAGTGAATrGrGrG-3’), and the resulting cDNA was amplified using 12 cycles (6,000 microglia input) or 18 cycles (1,000 microglia input) and KAPA HiFi Hot Start Mix (Roche, KK2602). After bead clean-up using 0.7x ratio with AMPure beads (Beckman Coulter, A63881), cDNA concentration was measured using the Qubit 1x dsDNA HS kit (Thermo Fisher Scientific, Q33231) and normalized to 0.4 ng/µl as input for library prep. 0.4 µl of each normalized sample was mixed with 1.2 µl of tagmentation mix containing Tn5 Tagmentation enzyme (Illumina, 20034198) and then incubated at 55 °C for 12 min. The reaction was stopped by quenching with 0.4 ul 0.1% SDS (Teknova, S0180) and followed by chilling the plate on ice for 10 min. 0.8 ul indexing primer (IDT) was added and amplified using 12 cycles with Kapa (non-hot Start, Kapa Biosystems, KK2102). Libraries were pooled and purified using two purification rounds with a ratio of 0.8x and 0.7x AMPure beads. Library quality was assessed using a Bioanalyzer (Agilent). PCR reactions were carried out on a 384-plate Thermal Cycler (BioRad). Pipetting steps were performed using the liquid-handling robots Dragonfly and Mosquito HV (SPT Labtech). Libraries were then sequenced using the Nextseq 550 high-output kit (Illumina, 20024907, paired end, 2 × 75 bp depth) or by Novogene on NovaSeq S4 (Illumina, paired end, 2 x 150 bp depth). Base calling, demultiplexing, and generation of FastQ files were conducted by Novogene.

For the middle-aged *Apoa1* KO mice rescue experiment, 11-month-old *Apoa1* KO mice were intravenously injected with ApoA-I protein (40 mg/kg) three times, with two days apart. One day before taking down the mice, Atto647N-labeled ApoA-I were injected into both wild-type mice and *Apoa1* KO mice. 20 hours later, enzymatic method was used as described above to isolate microglia, and 1,000 microglia from each group were sorted into 10x lysis buffer (Takara) and frozen at −80 °C. Total mRNA was transcribed into full-length complementary DNA using a SMART-Seq mRNA LP (Takara, 634768) according to the manufacturer’s instructions. cDNAs were validated using a Bioanalyzer (Agilent). Full-length cDNA (5 ng) was processed using a SMART-Seq Library Prep Kit (Takara, 634764) according to the manufacturer’s protocol. Library quality was assessed using a Bioanalyzer (Agilent). Libraries were sequenced by Novogene on NovaSeq S4 (Illumina, paired end, 2 x 150 bp depth). Base calling, demultiplexing, and generation of FastQ files were conducted by Novogene.

### Sort microglia for mass spectrometry proteomics

Twenty hours after injecting Atto647N-plasma into 3-months-old mice, plasma-positive microglia and plasma-negative microglia were isolated using ice-cold dounce method as described above. 12,000 microglia were sorted into RIPA buffer with Halt protease inhibitor cocktail (Thermo Fisher Scientific, 87785) per replicate from pooled mice.

For mass spectrometry sample preparation, as previously described^98^, the proteins were first precipitated by adding 4× volume of pre-cooled acetone (Honeywell). Then, the protein pellet was sequentially washed with pre-cooled acetone, pre-cooled ethanol, and then pre-cooled acetone again. The pellet was resuspended with UA buffer (8 M urea in 0.1 M Tris-HCl), then dithiothreitol (DTT, Sigma, 2 mM) was added before incubation at 30 °C for 1.5 hours. Iodoacetamide (IAA, Sigma, 10 mM) was added to the sample, and the sample was incubated and protected from light at 30 °C for 40 min. Afterwards, trypsin was added (final concentration 0.25 μg/μl) to the sample for overnight digestion. On the second day, the digestion was stopped by adding trifluoroacetic acid (TFA, Sigma, final concentration 0.4%). After salt depletion using C18 spin tips (Thermo Fisher Scientific), the samples were loaded onto the Bruker timsTOF Pro mass spectrometer following the manufacturer’s instructions. For Liquid chromatography (LC), the mobile phase A contains 98% MilliQ Water, 2% Acetonitrile with 0.1% formic acid, and the mobile phase B contains 100% Acetonitrile with 0.1% Formic acid. The ionoptiks 25 cm Aurora Series emitter column with CSI (25 cm x 75 µm ID, 1.6 µm C18) was used. For a detailed description of the timsTOF mass spectrometer, please see Zeng et al^98^.

Briefly, the Captive Spray Ion source initiates the generation of ions, which then enter the initial vacuum stage and accumulate in the front section of the dual Trapped Ion Mobility Spectrometry (TIMS) analyzer. The radial confinement of the ion cloud is maintained by a 300 Vpp RF potential. Following the initial accumulation, ions are transferred to the second section of the TIMS analyzer for ion mobility analysis. Both sections of the TIMS analyzer utilize an electrical field gradient (EFG) superimposed on the RF voltage. Consequently, the ions in the tunnel experience simultaneous dragging by the incoming gas flow from the source and retention by the EFG. Subsequently, the ions are released from the TIMS analyzer in accordance with their ion mobility for QTOF mass analysis. The dual TIMS setup enables 100% duty cycle operation by keeping accumulation and ramp times equivalent. Specifically, the accumulation and ramp times are set at 100 ms each, and mass spectra are recorded in the m/z range of 100–1700 using positive electrospray mode. Ion mobility scanning occurs from 0.85 to 1.30 Vs/cm². The comprehensive acquisition cycle of 0.53 s encompasses one full TIMS-MS scan and four Parallel Accumulation-Serial Fragmentation (PASEF) MS/MS scans. Calibration of the TIMS dimension is achieved linearly using three selected ions from the Agilent ESI LC/MS tuning mix [m/z, 1/K0: (622.0289, 0.9848 Vs cm^−2^), (922.0097, 1.1895 Vs cm^−2^), (1221,9906, 1.3820 Vs cm^−2^)] in positive mode.

The raw data was processed by PEAKS software (Version: X+). Protein searches were made against a Mus musculus reference proteome database downloaded from UniProt. The parent ion was 15 ppm, and the fragment ion was 0.05 Da. The protein FDR was set to 1%. We used spectral counting for the label-free quantification as described before^99^ and generated the normalized spectral abundance factor (NSAF) for comparisons. NSAF was calculated as the number of spectral counts (SpC) identifying a protein, divided by the protein’s length (L), divided by the sum of SpC/L for all proteins in the experiment. Finally, we multiplied the NSAF value by 10^6^. The two-tailed Student’s *t*-test was performed to determine the differentially expressed proteins between samples. The level of statistical significance was set at *P* < 0.05.

### Microglia sorting for metabolomics

Twenty hours after injecting Atto647N-plasma into 3-months-old mice, plasma-positive microglia and plasma-negative microglia were isolated using ice-cold dounce method as described above from hypothalamus. 50,000 microglia were sorted into 1% ultrapure BSA (Thermo Fisher Scientific, AM2618) in PBS per replicate from pooled mice. Cell pellet was washed once with mass-spec grade PBS, and then was resuspend in ice-cold 80% methanol in LC/MS water containing 500 nM isotope-labelled amino acids used as internal standards (Cambridge Isotope Laboratories). After vortexing for 10 min at 4 °C, samples were span down at max speed at 4°C for 15 min. The supernatant was transferred to a new tube, fully dried by Speedvac and stored at −80 °C until analyzed. On the day of analysis, the samples were reconstituted by 80% methanol, vortexed for 15 min at 4 °C and centrifuged at 17,000 g for 15 min. The supernatants were transferred to a new set of tubes, centrifuged again at 17,000 g for 15 min. The final supernatants were transferred to glass insert vials for LC/MS.

The characterization of polar metabolites was performed by liquid chromatography-mass spectrometry (LC/MS), employing a hydrophilic interaction liquid chromatography (HILIC) column coupled with an ID-X Tribrid mass spectrometer. Analytical methodologies utilized pooled samples for the acquisition of data-dependent tandem mass spectrometry (MS/MS) spectra. The subsequent unbiased differential analysis was conducted using the Compound Discoverer software, which utilizes databases from both local and web-based libraries. For the purpose of internal standardization, isotopically labelled amino acids were deployed. Detailed experimental procedures are detailed on protocol.io under the identifiers 10.17504/protocols.io.36wgqj3p3vk5/v1 and 10.17504/protocols.io.n2bvj83exgk5/v1^100,101^.

### Microglia sorting for lipidomics

Twenty hours after injecting Atto647N-plasma into 3-months-old mice, plasma-positive microglia and plasma-negative microglia were isolated using ice-cold dounce method as described above. 50,000 microglia were sorted into 1% fatty acid free BSA in PBS per replicate from pooled mice. Cell pellet was washed once with mass-spec grade PBS, and then was resuspend in 20 µl 80% mass-spec grade methanol. Samples were vortexed for 5 min at 4 °C and spin down. The supernatants were transferred to a tube with 500 µl chloroform:methanol (CHCl_3_/MeOH) at ratio of 2:1 (v/v) containing SPLASH LipidoMIX internal standard mix (Avanti). The samples were vortexed for 1 hour at 4°C, store at −80 °C.

On the next day, 100 µl 0.9% (w/v) NaCl were added, and samples were vortexed for 10 min at 4 °C. The mixture was centrifuged at 3,000 g for 5 min at 4 °C. The lower phase containing lipids was collected and dried using a SpeedVac. The samples were stored at −80°C. On the day of analysis, dried lipid extracts were reconstituted in 30 μl of acetonitrile (ACN):isopropyl alcohol:H_2_O at a ratio of 13:6:1 (v/v/v) and vortexed for 10 min at 4 °C. The samples were centrifuged for 15 min at 4 °C at the max speed, and transferred to glass insert vials for LC/MS.

Untargeted lipidomics involved the profiling of lipids through LC/MS utilizing an C18 column in conjunction with an ID-X tribrid mass spectrometer. The untargeted analysis of lipid species was carried out by LipidSearch. Normalization of the dataset was achieved by internal standardization using deuterium-labeled lipids. Comprehensive methodological details are accessible via protocol.io at the links 10.17504/protocols.io.5qpvor3dbv4o/v1 and 10.17504/protocols.io.3byl4jq6jlo5/v^100,101^.

### Single cell RNA-seq library preparation

For droplet-based scRNA-seq, twenty hours after injecting Atto647N-plasma into 3-month-old mice, plasma-positive microglia and plasma-negative microglia were isolated from hypothalamus, using ice-cold dounce method as described above, except that EDTA was removed from both MACS and FACS buffer. Approximately 20,000 microglia were sorted into receiving buffer (1% ultra BSA in PBS + RNAse inhibitor) and were loaded in the generation of GEMS (gel bead in emulsions). The library preparation was performed according to the Chromium Single Cell 3’ Reagents kit v3.1 user guide (10x Genomics) according to the manufacturer’s instructions. Library quality was assessed using a Bioanalyzer (Agilent). The libraries were sequenced and to an average depth of 60,000-80,000 reads per cell by Novogene on NovaSeq S4 (Illumina, paired end, 2 x 150 bp depth). Base calling, demultiplexing, and generation of FastQ files were conducted by Novogene.

For flow index sorting and plate-based scRNA-seq, cell lysis, first-strand synthesis and cDNA synthesis were performed using the Smart-seq-2 protocol as described previously in 384-well format^90,102^. In brief, microglia (CD45^low^ CD11b^+^ CD206^-^) were sorted using MA900 sorters (Sony) on the highest purity setting (‘Single cell modes’). Plasma-Atto647N fluorescence was recorded for each cell and corresponding sorted well. cDNA synthesis was performed using the Smart-seq2 protocol. After cDNA amplification (23 cycles), 0.16 ng cDNA was used to prepare libraries using Tn5 Tagmentation enzyme (Illumina, 20034198), following the manufacturer’s instructions. Library quality was assessed using a Bioanalyzer (Agilent). Pipetting steps were performed using the liquid-handling robots Dragonfly and Mosquito HV (SPT Labtech). Libraries were then sequenced using the Nextseq 550 high-output kit (Illumina, 20024907, paired end, 2 x 75 bp depth). Samples were sequenced at an average of 1 million reads per cell.

### Human iPSC acquisition and culture

The parental WTC11 (GM25256; Coriell) and WTC11-mEGFP (AICS-0036-006; Coriell) human iPSC lines were acquired through the Allen Cell Collection, available from Coriell Institute for Medical Research. The parental human iPSC line (ADRC 76) was generated by the University of California Alzheimer’s Disease Research Center (UCI-ADRC) from subject fibroblasts under approved Institutional Review Board and human Stem Cell Research Oversight protocols and provided by the Institute for Memory Impairments and Neurological Disorders. Non-integrating Sendai virus (CytoTune-iPS 2.0 Sendai Reprogramming Kit; A16517; Thermo Fisher Scientific) was used to perform reprogramming thereby avoiding any integration-induced effects. iPSCs were confirmed to be karyotypically normal by Array Comparative Genomic Hybridization (Cell Line Genetics), sterile, and pluripotent through Pluritest Array Analysis and trilineage *in vitro* differentiation.

All human iPSC lines were maintained on matrigel-coated plates (CB356238; Fisher scientific) according to the manufacturer’s specifications in TeSR-E8 (05990; STEMCELL Technologies) in 5% O_2_, 5% CO_2_ at 37°C. Cultured media was replenished every day with fresh medium. Cells were passaged every 5–7 d using ReLeSR (100-0484; STEMCELL Technologies) in TeSR-E8 media with 0.25 μM Thiazovivin (STEMCELL Technologies) according to the manufacturer’s specifications. CRISPR generation of CSF1R-G795A Homo knock-in (KI) iPSCs was performed following the published protocol by Chadarevian et al^45^.

### Differentiation of Hematopoietic progenitor cells (HPCs) and microglia (iMG) from iPSCs

HPCs and iPSC-derived microglia (iMG) were differentiated following the highly replicated protocol published by McQuade et al^57^. To begin HPC differentiation, iPSCs were passaged onto 6-well matrigel-coated plates in TeSR-E8 at a density of 80 colonies of 100 cells each per 35 mm well. On day 0, cells were transferred to Medium A from the STEMdiff Hematopoietic Kit (05310; STEMCELL Technologies). On day 3, cells were exposed to Medium B, and remained in Medium B for 7 additional days while small round HPCs began to lift off the colonies. On day 10, non-adherent CD43^+^ HPCs were collected by carefully removing medium and non-adherent cells with a serological pipette. At this point, HPCs were frozen in BamBanker freezing solution (Fisher scientific, NC9582225) for long-term storage. Cells used for HPC transplantation were later thawed in complete iMG medium (DMEM/F12 [11039-021; Thermo Fisher Scientific], 2× insulin-transferrin-selenite [41400045; Thermo Fisher Scientific], 2× B27 [17504044; Thermo Fisher Scientific], 0.5× N2 supplement [17502048; Thermo Fisher Scientific], 1× Glutamax [35050-061; Thermo Fisher Scientific], 1× non-essential amino acids [11140-050; Thermo Fisher Scientific], 400 mM monothioglycerol, and 5 mg/mL human insulin [I2643; Sigma] freshly supplemented with 100 ng/mL IL-34 [Cat #200-34; Peprotech], 50 ng/mL TGFβ1 [Cat #100-21; Peprotech], and 25 ng/mL M-CSF [Cat #300-25; Peprotech]) for 18–24 h to recover before being resuspended at 62,500 cells/µl in 1× DPBS (low Ca^2+^, low Mg^2+^) for transplantation. For iMG differentiation, HPCs were cultured on 6-well matrigel-coated plates in complete iMG medium for 28 days. Fresh complete iMG media was added every 48 h.

### iMG phagocytosis assays

Phagocytic activity was examined using the IncuCyte S3 Live-Cell Analysis System (Sartorius). ADRC76 WT iMG were plated at 50,000 cells/well (n=6 per condition) in complete iMG media on matrigel-coated 96-well plate. At noted t=0, 488-labeled fibrillized-Aβ(1-42) (fAβ; Anaspec) was added (2 µg/mL) to all treatment wells and ApoA-I protein (Athens Research &Technology, 16-16-120101-LEL) was added (100 µg/mL) to first treatment condition. No fAβ was added to the negative control group. At noted t=12 hr, ApoA-I protein was added (100 µg/mL) to second treatment condition. No ApoA-I was added to the fAβ control groups. During live imaging, four fields of view at 20x per well of phase and fluorescence were captured every hour in the IncuCyte S3 live cell. Using IncuCyte 2020B software, image masks for fluorescent signal (phagocytosis of 488-labeled fibrillized-Aβ) were normalized to total cell area (phase).

### Adult Intracranial Transplantation

All mouse surgeries and use were performed in strict accordance with approved National Institutes of Health (NIH) and institutional guidelines. Direct bilateral intracranial injections were performed as detailed in Hasselmann et al^103^. Briefly, M-CSF^h^ adult mice (2-month-old) were anesthetized under continuous isoflurane and secured to a stereotaxic frame (Kopf), and local anesthetic (Lidocaine 2%; Medline, 17478-711-31) was applied to the head before exposing the skull. Using a 30-gauge needle affixed to a 10-mL Hamilton syringe, mice received 2 μL of cell suspension in sterile 1× DPBS (14190144; Thermo Fisher Scientific) at 62,500 cells/μL at each injection site. HPC transplantation was conducted bilaterally into the lateral parietal association cortex and dorsal hippocampus at the following coordinates relative to bregma: anteroposterior, 2.06 mm; mediolateral, ± 1.75 mm; dorsoventral, 1.75 mm (hippocampus) and 0.95 mm (cortex). Cells were injected at a rate of 62,500 cells/30 s with 4 min diffusion time in between injections. The needle was cleaned with consecutive washes of DPBS, 70% (vol/vol) ethanol, and DPBS in between hemispheres and animals. Animals were allowed to recover on heating pads before being placed in their home cages and received 2 mg/mL Acetaminophen (Mapap; Major, 0904-7014-16) diluted in water for 10 days.

### CSF1Ri-mediated microglial replacement

CNS-wide microglia replacement was performed following previously published protocol by Chadarevian et al^45^. Briefly, PLX3397 (HY-16749) were obtained from MedChemExpress and formulated in AIN-76A standard chow by Research Diets Inc. at 290 mg/kg dose as noted. 2 weeks after HPC transplantation, adult mice were fed PLX-chow *ad libitum* for 2 months. Mice were then transitioned to standard chow for an additional 1 month before *in vivo* experimentation.

### xMG isolation from human microglia transplanted mice

Twenty hours after saline or plasma injection, mice were perfused with 1x PBS and brains were collected and micro dissected for the cerebellum, thalamus, hypothalamus, striatum, hippocampus, and cortex. Microdissected regions were immediately placed in 4 mL RPMI1640 on ice then transferred for xMG isolation as previously described by Chadarevian et al^44^. Briefly, micro dissected regions were manually homogenized using a 7 mL Dounce homogenizer by adding 4 mL of RPMI1640 and performing 10 strokes with the “A” pestle followed by 10 strokes with the “B” pestle. Samples were then run through 70-μM filters, presoaked in RPMI1640, and the filter was washed with 10 mL of RPMI1640. The samples were centrifuged, supernatant was discarded, and the pellets were resuspended in 30% Percoll overlaid with 2 mL of 1×DPBS centrifuged at 400×g for 20 min with acceleration and brake set to 0 to remove myelin. The myelin band and supernatant were discarded, and cell pellets were resuspended in 80 µL MACS buffer (0.5% BSA in 1×DPBS) + 20 µL Mouse Cell Removal beads (130-104-694; Miltenyi) and incubated at 4°C for 15 min. Magnetically labeled mouse cells were separated using LS columns and the Midi MACs separator (130-042-302; Miltenyi) while the unlabeled human cells were collected in the flow through. Human cells were then pelleted by centrifugation and resuspended for flow analysis.

### Primary rat microglia culture

Primary rat microglia were cultured in serum-free culture medium following a published protocol^104,105^. Briefly, single cell suspension was obtained from postnatal 14 (P14) rat brains using an ice-cold dounce (Wheaton) homogenize method. Myelin debris were removed firstly by Percoll spin (GE Health) and then by myelin removal beads (Miltenyi). Microglia were further isolated using CD11b microbeads and a column MACS separation system (Miltenyi). Purified microglia were resuspended and plated in microglial growth medium (MGM) at a density of 2 × 10^4^ cells/ml. Every 2 days, 50% of the medium was removed and equal volume of fresh medium was added. At day 10-12, functional assay was performed.

MGM was sterile-filtered and stored at 4°C for up to one month and was comprised of: DMEM/F12 containing 100 units/mL penicillin, 100 mg/mL streptomycin, 2 mM glutamine, 500 ng/ml N-acetyl cysteine, 10 µg/mL apo-transferrin, and 10 ng/mL sodium selenite (Sigma, 214485), human TGF-β2 (2 ng/mL, Peprotech, 100-35B), murine CSF1 (1 ng/mL, Peprotech, 315-02), heparan sulfate (1 µg/mL, Galen Laboratory Supplies), oleic acid (100 ng/mL, Cayman chemicals, 90260), gondoic acid (1 ng/mL, Cayman chemicals, 20606), and ovine wool cholesterol (1.5 µg/mL, Avanti Polar Lipids, 700000P).

### Biotin-tag enriched plasma proteomics

For *in vitro* Blood-brain Barrier (BBB) model, mouse endothelial cells (bEnd.3, ATCC, CRL-2299) were seeded at 2 × 10^5^ cells per 12 well transwell insert (3 μm pore size, Falcon 353181) and incubated for 3 days to reach full confluency. GSK-3 inhibitor BIO (2.5 μM, Sigma) was added to stimulate the Wnt/β-catenin pathways and increase electrical resistance^106^. The confluency was tested by classic tracer permeability assay using 10 kDa and 40 kDa fluorescein Dextran (Thermo Fisher Scientific).

Albumin and immunoglobulin (IgG) were depleted from dialyzed mouse plasma using an immunoaffinity Albumin & IgG depletion column (Sigma, PROTIA). Biotin-labeled IgG/Albumin depleted plasma was prepared as described above. 10% of biotin-labeled plasma (unfiltered plasma) was added to the upper chamber of the transwell seeded with the confluent endothelial cells. 4 hours later, BBB filtered plasma proteins were collected from the bottom chamber and were added to the primary microglia for 6 hours. In parallel, unfiltered plasma and unlabeled plasma were added to the primary microglia for 6 hours. Microglia were washed 3 times with 1x PBS and lysed by RIPA buffer with halt protease inhibitor cocktail (Thermo Fisher Scientific). n = 3 replicates per groups.

To enrich biotinylated proteins from samples and perform on-bead trypsin digestion, a published protocol was followed with minor modifications^107^. Briefly, pre-washed Pierce streptavidin magnetic beads (Thermo Fisher Scientific, 88817) were added to the microglia lysate and incubated at 4°C overnight with rotation. The beads were subsequently washed once with 1 mL of 50 mM Tris-HCl (pH 7.5), followed by two washes with 1mL 2 M urea in 50 mM Tris (pH 7.5) buffer. Then beads were incubated with 80 μL of 2 M urea in 50 mM Tris-HCl containing 1 mM DTT and 0.4 μg trypsin (Promega, V5073) at 37 °C for 1 hour while shaking at 1,000 r.p.m. After 1 hour, supernatant was removed and transferred to fresh tubes. Beads were washed twice with 60 μL of 2 M urea in 50 mM Tris-HCl (pH 7.5) buffer and the washes were combined with the on-bead digest supernatant.

Disulfide bonds in the eluate were reduced by adding DTT to a final concentration of 4 mM and incubated at 37 °C for 30 min with shaking at 1,000 r.p.m. The eluate was alkylated by adding iodoacetamide to a final concentration of 10 mM and incubated at 37 °C for 45 min in the dark while shaking at 1,000 r.p.m. An additional 0.5 μg of trypsin was added to the sample and digestion proceeded overnight at 37 °C with shaking at 700 r.p.m. The sample was acidified by adding formic acid (FA) so that the sample contains ∼1% (vol/vol) FA and at pH 3. UltraMicroSpin C18 columns (Nest Group, SUM SS18V) were used to desalt the samples following the manufacturer’s instructions. The peptide concentration was measured by quantitative peptide assay (Thermo Fisher Scientific, 23290).

### Mass spectrometry-based proteomics

LC-MS/MS data were acquired on a Q Exactive HF-X mass spectrometer (Thermo Fisher Scientific) equipped with a Nanospray Flex ion source (Thermo Fisher Scientific), active background ion reduction device (ABIRD) (ESI Source Solutions), butterfly portfolio heater (Phoenix S&T), and UltiMate 3000 RSLCnano chromatograph (Thermo Fisher Scientific) with a 50-cm PicoFrit Self-Pack Column with 75 µm inner diameter (New Objective) packed with 1.9-µm C18-coated beads with 120-Å pores (Reprosil-Pur C18-AQ, Dr. Maisch / ESI Source Solutions).

A 130-min trap-elute method was used for loading, separating, and eluting the peptides using a mixture of Buffer A (0.1% formic acid in HPLC-grade water) and Buffer B (0.1% formic acid in HPLC-grade acetonitrile). The column was heated to 50 °C. Peptides were loaded onto a trap cartridge for 5 min at 10 µl/min using 100% Buffer A and then loaded on to the column and separated at 300 nl/min using a multistep gradient: isocratic 4% Buffer B for 13 min, then a linear gradient to 30% Buffer B for 72 min, then a linear gradient to 40% Buffer B for 15 min, then a linear gradient to 96% Buffer B for 5 min, followed by 9 min of isocratic flow at 96% Buffer B, a 1-min linear gradient to 4% Buffer B, and then 10 min of isocratic flow at 4% Buffer B.

A 130 min data-dependent acquisition method was used. MS1 scans were conducted at 60,000 resolutions with an automatic gain control (AGC) target of 3×10^6^ and a maximum injection time (IT) of 20 ms with a scan range of 300–1650 Th in centroid mode. Features surpassing an intensity of 5.4×10^4^ charges and having an assigned charge state of 2–5 were selected for MS2, with peptide-like (“peptide match” option) features preferred. The dynamic exclusion duration was 45 s and isotopes were excluded. Up to 15 features per MS1 were selected per MS2. Precursors were selected using a 1.4-Th isolation window and fragmented using higher-energy collision-induced dissociation at 28% normalized collision energy. MS2 scans were conducted at 15,000 resolutions with an AGC target of 1×10^5^ and a maximum IT of 54 ms with a scan range of 200–2000 Th in centroid mode. A lock mass of 445.12003 Th was used to maintain mass accuracy.

### Proteomic data analysis

Raw data were processed using MaxQuant (version 1.6.10.43)^108^ and quantitative comparisons were performed using Perseus (version 1.6.2.3)^109^. Peptide spectral matches (PSMs) were made against a mouse reference proteome database downloaded from UniProt using a target-decoy Andromeda search. Methionine Oxidation and protein N-terminal acetylation were specified as variable modifications, and carbamidomethylation of cysteine was specified as a fixed modification. A precursor ion search tolerance of 20 ppm and product ion mass tolerance of 20 ppm were used for searches. Both unique and razor peptides were used for quantification. Cleavage specificity was set to trypsin/P, with up to two missed cleavages allowed. Results were filtered to a 1% false discovery rate (FDR) at both the peptide and protein levels. Proteins were quantified and normalized using MaxLFQ^110^ with a label-free quantification (LFQ) minimum ratio count of 1. The “second peptides” and “match between runs” features were enabled.

For quantitative comparisons, protein intensity values were first log_2_-transformed, proteins with at least two measured values in at least one group per condition were retained, and missing values were imputed from a normal distribution with a width of 0.3 and a down shift value of 1.5 using Perseus. Principal component analysis (PCA) analysis was performed in Perseus using the Benjamini-Hochberg FDR with a cut-off of 0.05. Significance calculations for pairwise comparisons were performed using a two-tailed *t*-test (*P* value < 0.05).

### ApoA-I uptake *in vitro* assay

Endothelial cells (bEnd3) were maintained in DMEM + 10% FBS (high glucose, pyruvate, Thermo Fisher Scientific). Before the uptake assay, culture medium was changed to Opti-MEM (Thermo Fisher Scientific) to reduce serum. Primary microglia were maintained in serum-free medium as described above. Blocking antibody against SR-BI (Novus, NB400-113, 1:100) or control IgG (Cell signaling technology, 2729s) was added to primary microglia (DIV10-12) or endothelial cells. 3 hours later, lipid free version of Atto647N-ApoA-I protein (5 µg/ml) was added for another 3 hours. After that, cells were digested and analyzed by flow cytometry to detect the level of ApoA-I uptake.

### Primary microglia with LPS stress

Primary microglia were cultured in the serum-free culture medium as described in the “primary microglia culture” section. On DIV12, lipopolysaccharide (LPS, Sigma L4391, 100 ng/ml) was added for 20 hours with or without ApoA-I protein (100 µg/ml). After 1x PBS wash, cells were lysed in the RLT buffer with beta-mercaptoethanol (beta-ME), and the RNA was isolated using RNeasy Plus Micro kit (Qiagen, 74034). Library preparations and bulk RNA-seq were done by Novogene with 30 million reads per sample.

### Bulk RNA-seq data analysis

Unless otherwise noted, raw sequencing files were demultiplexed with bcl2fastq to generate fastq files and were trimmed to remove adaptor contamination and poor-quality reads using Trim Galore!. Trimmed reads were aligned using STAR, multi-mapped reads were filtered, and gene counts were made using HTSEQ. Data visualization and analysis were performed using Seq-Monk v.1.48.0 and R version 4.1.2 using the following Bioconductor packages: DESeq2 v.1.34^111^, topGO v.2.44. The ‘elim’ algorithm were used in topGO. Unless stated otherwise, the set of expressed genes (RPKM of ≥ 1 in at least 3 samples) was used as background for functional enrichment analyses. *P* values were adjusted for multiple testing and significant DEGs were defined by a cutoff of adjusted *P* < 0.05.

For *Apoa1* KO mice rescue bulk RNAseq dataset, sixteen single-end fastq files were generated, consisting of four replicates each for the following groups: *Apoa1* knockout mice-ApoA-I positive microglia (KO-A1-Pos), *Apoa1* knockout mice-ApoA-I negative microglia (KO-A1-Neg), wild-type mice-ApoA-I positive microglia (WT-A1-Pos), and wild-type mice-ApoA-I negative microglia (WT-A1-Neg). For the primary microglia with LPS stress dataset, sixteen double-end fastq files were generated, consisting of four replicates each for the following groups: Con, LPS, ApoA-I, and LPS+ApoA-I.

Adapter sequences and low-quality base calls (Phred score < 30 at ends) were trimmed using Cutadapt v1.8.1^112^. Trimmed reads were aligned to the reference genome using STAR 2.7.10b^113^ with comprehensive gene annotations from GENCODE release vM30. Gene-level transcript abundances were quantified with RSEM v1.3.1^114^. RNA-Seq read counts and Transcripts Per Million (TPM) values were obtained from RSEM output and imported into R software for further analysis. Genes were considered expressed if their TPM exceeded 0.5 in at least 80% of samples within each group. Only expressed genes were retained for subsequent analyses. Principal component analysis (PCA) was performed using all expressed genes. Z-scores were calculated for the TPM values across all samples.

Differential gene expression (DGE) was assessed using DESeq2^111^. Genes with a *P*-value < 0.05 were considered differentially expressed and were retained for downstream enrichment analysis for *Apoa1* KO mice rescue dataset and adjust *P*-value < 0.05 were considered differentially expressed and were retained for downstream enrichment analysis for primary microglia with LPS stress dataset. Enrichment analysis for Gene Ontology (GO) biological processes pathways was performed using topGO v2.44. The background gene set for this analysis was restricted to the set of expressed genes, ensuring relevant context for enrichment assessments. The top 200 relevant pathways were calculated with node size = 10.

### scRNA-seq data analysis

10x Single-cell RNA-seq binary cell call (BCL) files obtained from Novogene were converted to fastq file using the mkfastq function from Cell Ranger^115^ version 3.0.1 with the default parameters. Fastq files were aligned to mm10 (Ensembl 93) reference transcriptome, and the count function of the Cell Ranger with the default parameters were used to generate gene expression matrices. R version 4.1.2 and Seurat^116^ package version 4.3 were used to perform all downstream analysis.

After checking the QC metrics (nCount_RNA, nFeature_RNA (1,000-5,000), percent.mitochondrial < 10%), the data was further filtered to remove low quality cells. SCTransform normalization^117^ from Seurat was used to normalize, find variable genes, and scale the dataset. Principal component analysis was done by using the identified list of variable genes to determine the dimensionality of the data. Clustering was done using the FindNeighbors (dims =1:8) and FindClusters function (resolution = 0.25) and assignment of cluster identity was done using the FindAllMarkers function. Once clusters were assigned and UMAP was generated using Seurat’s RunUMAP function, the data was ready for differential expressed gene (DEG) analysis. The Seurat function FindMarkers function was used with the MAST test to produce a DEG list. Creation of microglial identity scores and *ex vivo* activation scores were executed using AddModuleScore function in Seurat. Score were calculated for microglial identity and *ex vivo* activation based on published gene lists^39^ (Table S7).

For the smart-seq2 dataset, raw sequencing files in blc format were demultiplexed with bcl2fastq to generate fastq files and were trimmed to remove adaptor contamination and poor-quality reads using Trim Galore!. Trimmed reads were aligned using STAR (v.2.5.3a) GeneCounts function, multi-mapped reads were filtered, and gene expression matrices were generated using SeqMonk. Seurat^116^ package was used to perform all downstream analysis. After checking the QC metrics (nCount_RNA, nFeature_RNA, percent.mitochondrial), the data was further filtered to remove low quality cells (200 ≤ nFeature ≤ 5000; percent.Mt ≤ 5). Data was integrated using Seurat’s built-in SCTranform, followed by embedding (top 20 PCs) and k-means clustering (top 20 PCs; resolution = 0.4).

During the index sorting step, the intensity of plasma signal within the microglia was recorded quantitatively allowing a separation of microglia into three groups: with no plasma signal, low plasma signal, and high plasma signal. DEG was performed in Seurat using the FindMarkers() function.

Plasma-positive gene signature for plasma score analysis is generated by selecting the top 96 most differentially expressed genes (upregulated) comparing PPM versus PNM ranked by adjust *P*-value (Table S7). Gene signatures are used in this study to quantify the expression of a gene set, thus representing the aggregated expression of multiple genes in a given transcriptome (here: list of plasma-positive gene signature in the microglia scRNA-seq data). The resulting value is defined as a score.

We generated signatures and quantified scores using the VISION (v.3.0.1) package as reported previously^38^. Notably, VISION z-normalizes signature scores with random gene signatures to account for global sample-level metrics (such as total number of counts/UMIs). In addition to the two datasets that we generated in our current study (the 10x single cell dataset and the smart-seq2 single-cell dataset), we also extended the application of the plasma score to two published mouse microglia datasets^29,42^ and two published human microglia datasets^43,89^. In the Moffit dataset^42^, microglia were isolated from hypothalamic region, given that our mouse data revealed the highest plasma uptake in hypothalamus. We putatively selected the top 10% of high-score microglia and the bottom 10% of low-score microglia for subsequent comparison. For the other three datasets, where microglia were isolated from either the whole brain, temporal neocortex^43^, or prefrontal cortex region^89^, we similarly putatively selected the top 5% of high-score cells and the bottom 5% of low-score cells for comparative analysis. Plot and *P*-value generated from VlnPlot function. DEG analysis was conducted on microglia with the highest and lowest VISION-calculated signature scores using FindMarkers function in Seurat.

### Statistics and reproducibility

Statistical analyses were performed using the software GraphPad Prism 10.2.2 (GraphPad Software). The DESeq2 package in R was used for bulk RNA-seq analysis. The Seurat package version 4.3 in R was used for scRNA-seq sequencing data. Statistical tests used, number of replicates and *P* values for each experiment are described in figures and figure legends.

Sample sizes were chosen based on prior literature using similar experimental paradigms. In addition, sample sizes were determined to be adequate based on the magnitude and consistency of measurable differences between groups. Box plots show 25–75th percentiles and the median, and the whiskers are created by Tukey method. Violin plot midlines indicate the median. For all statistical tests shown, ns: non-significant, \**P* < 0.05, \*\**P* < 0.01, \*\*\**P* < 0.001, \*\*\*\**P* < 0.0001.

## Data and code availability

Data availability: Single-cell data and bulk RNA-seq datasets are available through Gene Expression Omnibus under accession code GSE263833, GSE264113, GSE264105, and GSE264111, and GSE264110.

The mass spectrometry proteomics data have been deposited to the ProteomeXchange Consortium via the PRIDE^118^ partner repository under dataset identifier PXD051375 for timsTOF proteomic data of plasma-positive and plasma-negative microglia (reviewer account details: username; password: Pgkrkiu6), and PXD051374 for proteomics data of biotin-tagged plasma uptake in primary microglia (reviewer account details: username; password: j74VBi0g).

Code availability: R code used to analyze single cell datasets is available on GitHub at (https://github.com/twclab/PPM).

## Supplemental Table

**Table S1**

Bulk RNA-seq data of plasma-positive microglia (PPM) and plasma-negative microglia (PNM) isolated from the hypothalamus of young mice injected with NHS-Atto647N labeled plasma proteins.

**Table S2**

Full Gene Ontology (GO) enrichment list of plasma-positive microglia (PPM) and plasma-negative microglia (PNM) from the hypothalamus of young mice injected with NHS-Atto647N labeled plasma proteins based on all the significantly differential expressed genes (*P* adjust < 0.05).

**Table S3**

timsTOF proteomic data of plasma-positive microglia (PPM) and plasma-negative microglia (PNM) isolated from the hypothalamus of young mice injected with NHS-Atto647N labeled plasma proteins.

**Table S4**

Full Gene Ontology (GO) enrichment list of plasma-positive microglia (PPM) and plasma-negative microglia (PNM) from the hypothalamus of young mice injected with NHS-Atto647N labeled plasma proteins based on all the significantly differential enriched proteins from timsTOF data (*P* adjust < 0.05).

**Table S5**

Metabolomic data of plasma-positive microglia (PPM) and plasma-negative microglia (PNM) isolated from the hypothalamus of young mice injected with NHS-Atto647N labeled plasma proteins.

**Table S6**

Lipidomic data of plasma-positive microglia (PPM) and plasma-negative microglia (PNM) isolated from the hypothalamus of young mice injected with NHS-Atto647N labeled plasma proteins.

**Table S7**

Microglial gene signature lists, including *ex vivo* activation score, microglia identity score, Plasma score (mouse), Plasma score (human), and 3 randomly selected 100 genes lists.

**Table S8**

Differential expression gene (DEG) list of Plasma score-high and Plasma-score low cells from Moffit, Olah, Hammond and Lau datasets (adjust *P* value < 0.05).

**Table S9**

Proteomic data of *in vitro* biotin-plasma enrichment assay (Filtered plasma versus Control group). n =3 replicates per group.

**Table S10**

Full Gene Ontology (GO) enrichment list of KO-A1-Pos groups versus KO-A1-Neg group from ApoA-I rescue experiment based on all the significantly differential expressed genes.

**Table S11**

Unique and overlapped gene list of plasma signature and disease-associated microglia (DAM) signature.

